# Erucamide regulates retinal neurovascular crosstalk

**DOI:** 10.1101/2025.09.02.673793

**Authors:** Guoqin Wei, Shreyosree Chatterjee, Qinglin Yang, Sanahan Vijayakumar, Daisuke Ogasawara, Sarah Giles, Peter Westenskow, Junhua Wang, Ruhan Fan, Helena Pham, Edith Aguilar, Jacob Robinson, Ayumi Usui-Ouchi, Roberto Bonelli, Kevin Eade, Gary Siuzdak, Benjamin Cravatt, Michael J. Sailor, Dale Boger, Martin Friedlander

## Abstract

Neurovasculoglial crosstalk is critical in establishing and maintaining a functional neurovascular unit. Breakdown in the unit is central to many neurodegenerative disorders of the CNS of which the retina is a component. A growing literature indicated that primary fatty acid amides (PFAMs) can regulate this crosstalk between vasculature and neuronal tissues. In this study we describe a central role for erucamide, a 22:1 mono-unsaturated omega-9 fatty acid amide, in degenerating retinal tissues. Using high-resolution global mass spectrometry-based metabolomics, we cataloged metabolites in murine models of retinal degeneration and show that while PFAMs, in general, are highly dysregulated, erucamide is the one most significantly diminished during photoreceptor atrophy. Using rodent models of retinal degeneration and novel organosilane-modified porous silicon nanoparticles (pSiNPs) for the *in vivo* delivery of erucamide, we demonstrate that erucamide activates CD11b+ myeloid cells, leading to the upregulation of angiogenic and neurotrophic cytokines that stabilize retinal degeneration. We identified TMEM19 as a novel binding protein for erucamide that is crucial for human iPSC-derived macrophage precursor cells activation and subsequent neurotrophic and angiogenic factor production. These findings reveal a previously unknown PFAM pathway that is modulated during retinal degenerative diseases, demonstrating that erucamide or functional analogues and their action through TMEM19 may be useful as a therapeutic alternative to neuroprotective and stem cell-based approaches for the treatment of retinal degenerative diseases.

## Main

The vast majority of diseases that lead to catastrophic loss of vision do so as a result of breakdown in the neurovascular/glial unit of the retina. Retinal pigmented epithelial (RPE) cell malfunction and subsequent photoreceptor degeneration can lead to vision loss, exemplified by conditions such as age-related macular degeneration (AMD). Other significant causes of vision loss are inherited retinal degenerations such as retinitis pigmentosa (RP) ^1^ and neurovascular retinal diseases such as diabetic retinopathy (DR).^2^ These diseases either do not have a clear genetic basis (e.g., DR) or are genetically heterogeneous (e.g., RP) ^3^ . While these diseases have different underlying causes, they consistently result in similar pathologies with functional loss of the interconnected vasculature, glia, photoreceptors, and RPE, thus leading to vascular anomalies and photoreceptor death.

In the neurosensory retina, photoreceptors convert light into electrical signals through a collaboration between neural retina, the RPE, Bruch’s membrane, and the choriocapillaris. This phototransduction process begins in the photosensitive outer segments of rod and cone photoreceptors. The RPE plays a critical role in this process by interdigitating with the photoreceptor outer segments where it, among other things, phagocytoses shed outer segments and recycles isomerized opsins. It’s functions are necessary for the activity and survival of photoreceptors.^4^ Tightly controlled lipid homeostasis is crucial to regulate the turnover between photoreceptors and RPE cells, and to maintain their cellular functions.^5^

In the retina, specific fatty acid amides play significant roles in maintaining retinal health and function. For example, ceramides are key players in various forms of photoreceptor cell death and are potential drug targets for retinal degenerative diseases^6^. Anandamide, also known as N-arachidonoylethanolamide, is a well-characterized neurotransmitter that regulates the endocannabinoid signaling pathway and can modulate visual signal processing by interacting with cannabinoid receptors in the retina^7^. However, little is known about the specific roles of PFAMs in the retina. As a direct extension of the central nervous system, the retina shares many characteristics with neurons of the brain. A better understanding of the biological and pathological functions of PFAMs in the retina will likely also shed light on their roles in the brain and other neuronal tissues.

Primary fatty acid amides (PFAMs)^8, 9^ represent a family of endogenous chemical messengers and signaling molecules in the mammalian nervous system. Structurally, PFAMs consist of a long hydrocarbon chain with an amide group (–CONH_2_) at one end. PFAMs like palmitamide, oleamide and the other three PFAMs were first discovered in 1989 in luteal phase plasma^10^. Although their biological functions were initially unclear, interest increased when oleamide was identified as an endogenous sleep-inducing molecule in mammalian cerebrospinal fluid^8^. Another PFAM, erucamide, was detected in the plasma and cerebrospinal fluid of pig brain^11^ and was first isolated from bovine mesentery. It was reported to be pro-angiogenic over 34 years ago^12, 13^, though its complete biological function remains unclear.

Given the critical role of PFAMs in the nervous system and the known importance of other fatty acids in the retina, we undertook an untargeted metabolomic and lipidomic screen of normal and degenerating retinas. We found that the most profound changes in the retinal lipid profile occurred with a PFAM, erucamide. We have identified erucamide’s target cells, binding proteins, and its neurotrophic mechanism of action. To address the challenge of *in vivo* delivery of the hydrophobic erucamide molecule, we utilized organosilane-modified porous silicon nanoparticles (pSiNPs). Intravitreal and subretinal injections of erucamide-NPs led to widespread activation of GS-Lectin positive cells, which were identified as CD11b+ myeloid cells. This activation was accompanied by the upregulation of angiogenic cytokines and growth factors, suggesting a novel therapeutic potential for erucamide in retinal diseases.

## Results

Using high-resolution global mass spectrometry-based metabolomics, we cataloged the metabolites dysregulated during retinal degeneration in an animal model of RPE-mediated retinal degeneration, the Royal College of Surgeons (RCS) rat. Dysregulated features were identified using our data processing technology (XCMS)^14^ combined with a database we developed (METLIN)^15^(Fig. 1a). Of all the untargeted metabolites, PFAMs emerged as one of the most severely dysregulated classes of metabolites in degenerating retinas. Of these, erucamide (22:1, n-9; C_22_H_43_NO; Fig. 1d), a 22:1 mono-unsaturated omega-9 fatty acid amide whose functions are not completely understood, is the most abundant PFAM in the eyes from wild-type rats. However, it is significantly reduced in the RCS rat model of photoreceptor atrophy (Fig. 1a, b and Extended Data Fig. 1a). Erucamide’s level is also largely decreased in the RD10 mouse, a rodent retinitis pigmentosa model of photoreceptor loss (Fig. 1c), suggesting that it may play a role in photoreceptor maintenance.

**Fig. 1.**
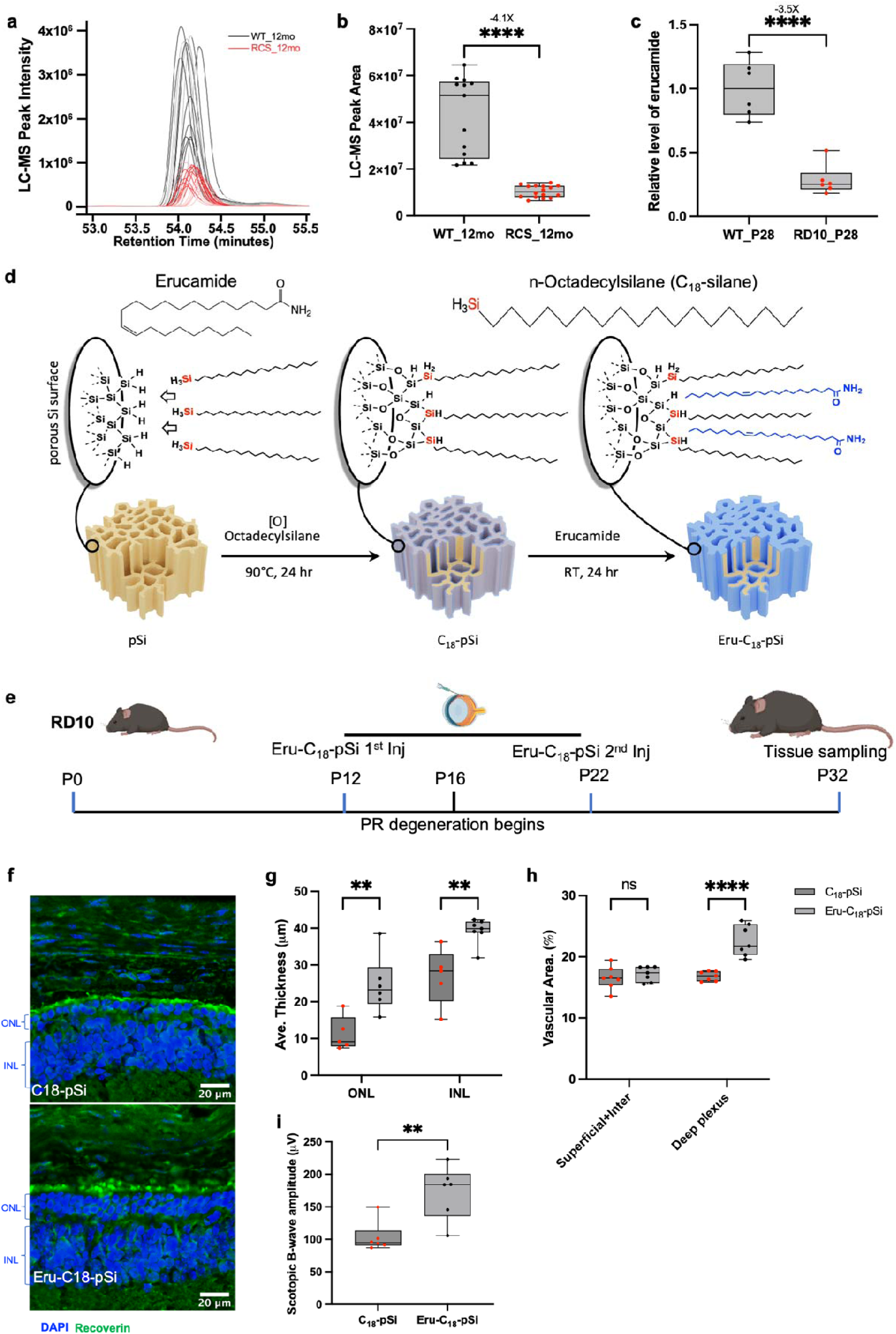
Erucamide is neurotrophic. **a**,**b**, Erucamide is highly dysregulated in one year old RCS eyes (n>13; p<0.0001). **c**, Erucamide level is attenuated in retinal degeneration mouse model RD10, 28.4% ± 0.1 (n=6; p<0.0001). **d**, Schematic illustration of the silane functionalization to the pSiNPs as well as erucamide loading to the C_18_-pSiNPs. **e**, Schematic illustration of Eru-C_18_-pSiNPs (214ng) intravitreal injection in RD10 mice. **f**, Retinas from P32 RD10 mice. Photoreceptors were immunolabeled green (recoverin). **g**, Effects of erucamide injections quantified in f (n=6; Outer nuclear layer (ONL): Eru vs. vehicle, p= 0.0033. Inner nuclear layer (INL): Eru vs. vehicle, p= 0.0047. **h**, Vascular density quantification of retinal vessels of superficial + intermediate, and deep plexus (SP+IP, and DP). n=7, SP+IP: Eru vs. vehicle, p= 0.7538. DP: p<0.0001. **i**, Electroretinographic (ERG) measurements of scotopic B-wave responses to a flash in dark-adapted RD mice injected with Eru or vehicle at P32 (n=6 per group). Eru vs. vehicle, p= 0.0048 at 50 cd*s/m2 intensity.

Erucamide is present in the central nervous system of many species including humans^8, 13, 16^ and has been shown to regulate angiogenesis and water balance in skeletal muscle and the intestine.^12, 13, 16^ However, its function in the central nervous system, biosynthetic pathway, site of action, and potential signaling cascades are all unknown. As erucamide is highly hydrophobic, direct retinal injection is not possible as it aggregates locally and fails to disperse within the retina. Consequently, we generated functionalized porous silicon nanoparticles (pSiNPs) for the delivery of erucamide both *in vivo* and in cultured tissues.

PSiNPs are biocompatible, showing low toxicity in multiple cell types (Extended Data Fig. 1f), and can biodegrade into silicic acid end products that are readily excreted from the body^17^. In addition, their tunable pore size and their readily functionalized surface make them amenable to the loading of a wide range of small-molecule and biologic therapeutics. As a result, pSiNPs have been used extensively to deliver drugs to sensitive and hard-to-reach tissues^18^, including the delivery of small-molecule drugs to the retina^19^. Chemical surface functionalization of pSiNPs with n-octadecylsilane (H_3_Si (CH_2_)_17_CH_3_, referred to here as C_18,_ Fig. 1e) imparts a hydrophobic character^20^ to the mesopores (10-15 nm), were thus chosen for loading of erucamide (Fig. 1d). The nanoparticles maintained desirable sizes (∼200 nm) and zeta potential (−33 mV) after erucamide loading, as shown by the TEM images, dynamic light scattering (DLS), and ζ-potential measurements (Extended Data Fig. 1a-c). The loading of erucamide was confirmed by the attenuated total reflectance Fourier transform infrared (ATR-FTIR) spectrum of the erucamide-loaded C_18_-pSiNPs (Eru-C_18_-pSiNPs), which displayed the characteristic C=O (at 1690 cm^-1^) and N-H (at 3310 cm^-1^) stretching bands of erucamide. The C-H scissor and stretching modes (at 1465 cm^-1^ and 2849 cm^-1^, respectively) are assigned to both erucamide and the C_18_ surface modification^21, 22^, while a band at 1080 cm^-1^ is assigned to the Si-O-Si stretching modes from silicon oxides on the surface of the pSiNPs. (Extended Data Fig. 1d). Thermogravimetric analysis (TGA) measurements indicated that the particles were composed of approximately 6% (wt.%) erucamide, based on the mass change upon oxidative combustion at 800 °C (Extended Data Fig. 1e).

Next, we investigated the potential therapeutic function of erucamide in the mammalian retina. RD10 mice were used in these experiments since vasodegeneration and photoreceptor degeneration starts at postnatal day 16-20 (P16-P20) and occurs in a stereotypical temporal sequence,^23^ and erucamide levels are spontaneously attenuated by P12 (Extended Data Fig. 1b). We injected 214 ng of erucamide into the subretinal space of RD10 at P12 and P22 through Eru-C_18_-pSiNPs (Fig. 1e). We observed that the extent of photoreceptor atrophy was significantly reduced in RD10 mice at P32 with erucamide. The number of surviving rows of photoreceptors in the eyes subretinally injected with Eru-C_18_-pSiNPs was significantly greater than the control eyes (injected with C_18_-pSiNPs) at P32 (6.4+-2.1 vs 1.8+-0.7, Fig. 1f). Similarly, the thicknesses of both the outer nuclear layer (ONL) and the inner nuclear layer (INL) of photoreceptors were rescued by erucamide. (Fig. 1g). Additionally, erucamide significantly protected the intraretinal deep plexus vasculature from vaso-obliteration compared to vehicle-injected controls (Fig. 1h). To further explore the importance of erucamide for retinal function, we used electroretinography to compare light responsiveness in the two groups of injected RD10 mice and observed significant rescue in rod-driven pathways in Eru-C_18_-pSiNPs injected mice (Fig. 1i), while cone-driven pathways remained unchanged (Extended Data Fig1. c-f). Taken together, these data showed that erucamide rescued retinal atrophy both morphologically and functionally in RD10 mice.

To evaluate its effects on healthy ocular tissue and to determine the cellular and biological target of erucamide, it was intravitreally administered into wild-type (WT) retinas. Eru-C_18_-pSiNP injected mice showed an infiltration of GS-Lectin+ cells at three days post injection, while retinas injected with C_18_-pSiNPs maintained a normal appearance similar to the non-injected control group (Fig. 2a). Further staining with Collagen IV (ColIV)^24^ and CD11b^25^ revealed that the GS-Lectin+ active signals were CD11b+ myeloid cells but not ColIV+ endothelial cells (Fig. 2b). These results suggest that erucamide might directly target CD11b+ myeloid cells in the retina. We synthesized a fluorescent dye (BODIPY) labelled erucamide (^BDP^Erucamide) (Fig. 2c) to directly observe the cellular targets. Strikingly, all ^BDP^Erucamide was taken up by CD11b+ cells, providing additional evidence that CD11b+ cells are the direct target of erucamide in the retinas (Fig. 2d). We further observed that subretinal injections of Eru-C_18_-pSiNP activated CD11b+ cells in superficial, intermediate, and deep vascular plexus layers throughout the retinas (Extended Data Video. 1).

**Fig. 2.**
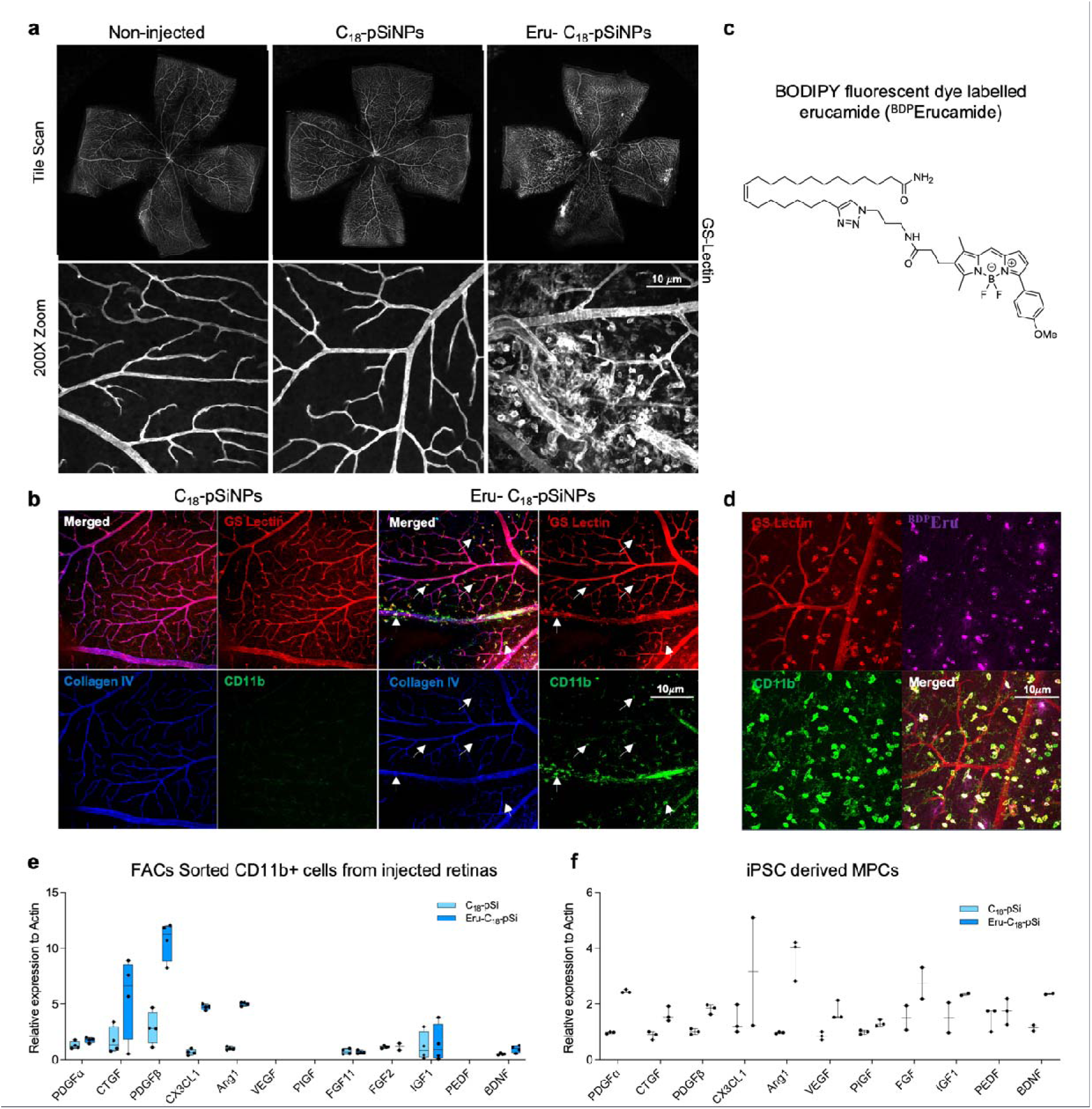
Erucamide delivered by pSiNPs targets CD11b+ cells in WT retinas. **a**, Eru-C_18_-pSiNP (214ng) injecte retinas exhibit increasing GS-Lectin positive active signals compared to vehicle treated and non-injected eyes. **b**, The GS-Lectin positive active signals are CD11b+ MPCs cells (pointed by white arrows). **c**, Structure of BODIPY fluorescent dye labelled erucamide (^BDP^Erucamide). **d**, ^BDP^Erucamide (214ng) was taken up by CD11b+ cells after injection. **e**,**f**, Angiogenic and neurotrophic factors are upregulated in FACs-sorted CD11b+ cells (**e**) from injecte retinas (214 ng erucamide, n=4) and iPSC MPCs (**f**) (0.64 µM erucamide, n=3) by erucamide treatment.

To further explore the potential effects of erucamide on the target cells, we isolated CD11b+ cells by flow cytometry and evaluated the expression of pro-angiogenic genes within the cells by qPCR. We found that the expression of PDGFα, CTGF, PDGFβ, CX3CL1, Ang1 and BDNF was higher in CD11b+ cells of erucamide-treated animals (Fig. 2e). Furthermore, the findings translate to human tissue. Treatment of human iPSC-derived macrophage precursor cells (MPCs)^26^ with erucamide led to significantly higher levels of PDGFα, VEGF, FGF2, IGF1 and BDNF compared to controls (Fig. 2f). Taken together, these data suggest that CD11b+ myeloid cells are the primary target of erucamide both *in vivo* and *in vitro* resulting in pro-angiogenic and neurotrophic factors.

These results raised the question of whether there is a specific receptor or binding protein for erucamide at the cellular level. We used a photoaffinity labeling (PAL) method^27^ to further explore and define the potential binding proteins of erucamide in a human microglia cell line, HMC3 cells. As shown in Fig. 3, we generated ‘fully functionalized’ lipid probes of erucamide (Eru.da) and the closely related PFAMs, which are also naturally present in the retina (oleamide and palmitamide, Ole.da and Pal.da), as control probes^27^. A photoreactive element (diazirine group) was introduced into the carbon chain of erucamide, oleamide, and palmitamide to convert reversible small molecular-protein interactions into stable, covalent adducts upon UV light irradiation. Introduction of a terminal alkyne handle allows late-stage conjugation to rhodamine or biotin tags by click chemistry (Fig. 3a, b). Erucamide, oleamide and palmitamide probes showed overlapping but distinct protein interaction profiles in HMC3 cells (Extended Data Fig. 3a). Following photoaffinity labeling, biotin conjugation and affinity enrichment, multiplexed proteomics of all three probes provided a unique list of erucamide-specific binding proteins that are involved in specific biological processes like lipid metabolism, signaling, and cellular processes. However, a significant number of these proteins were uncategorized as membrane traffic-related proteins (Extended Data Fig. 3b, c). From the list of proteins identified, a transmembrane protein, TMEM19 was the most likely candidate to be an erucamide binding protein (Fig. 3c) since it was the mostly enriched protein in PAL of erucamide diazirine probe compared to control probes. We confirmed the interaction between erucamide and TMEM19 by overexpressing TMEM19 in HEK293T cells and observed that Over-expressed TMEM19 is prominently labeled by erucamide probe compared to control probes (Fig. 3d). These data indicate that TMEM19 interacts with erucamide and is one of the mechanisms by which erucamide impacts cell function.

**Fig. 3.**
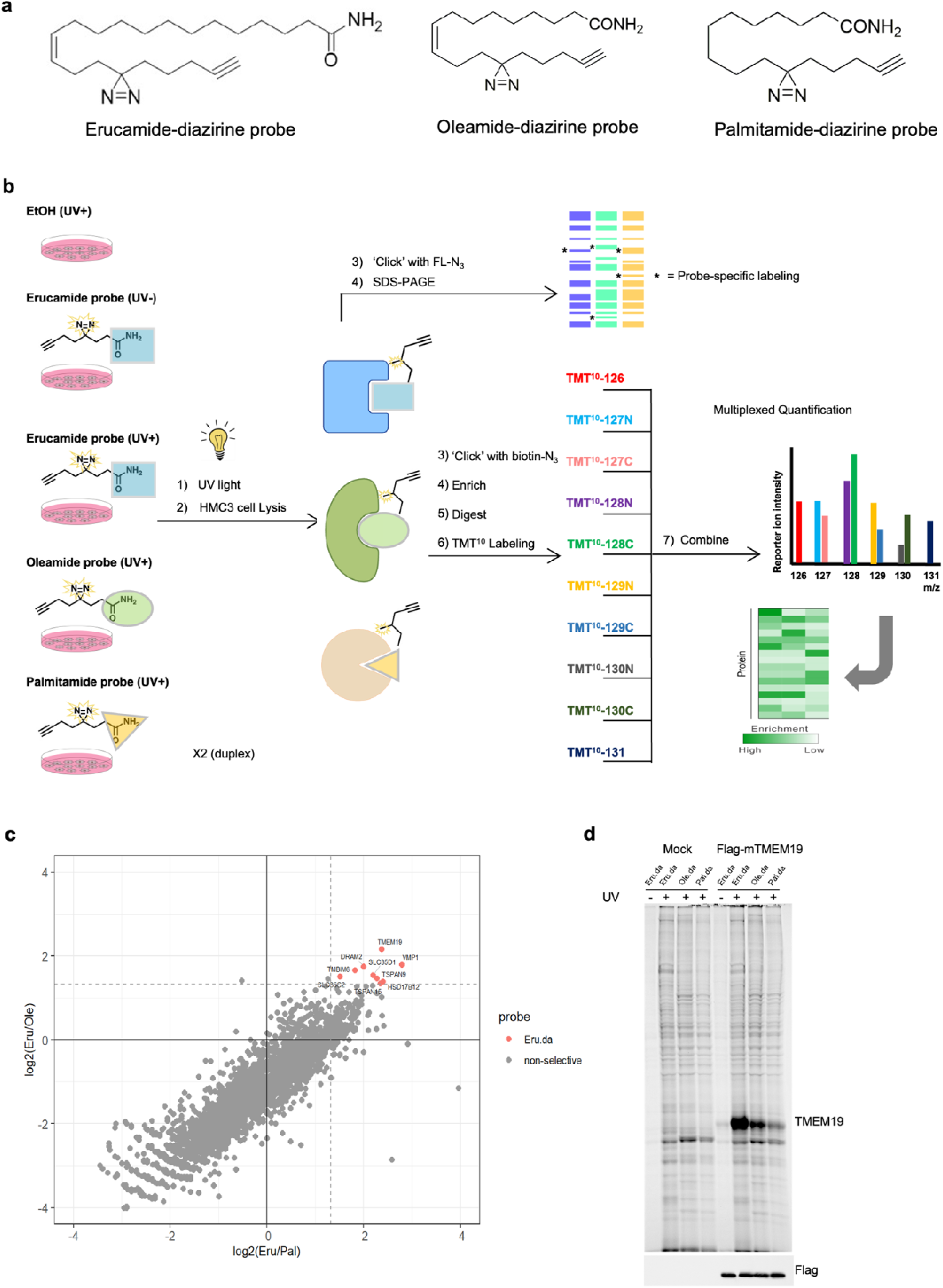
Identifying the binding protein of Erucamide in microglia by photoaffinity labeling (PAL). **a**, ‘Fully functionalized’ lipid probes of erucamide, oleamide and palmitamide possessing photoreactive diazirine and alkyn handles. **b**, Chemoproteomics experiment workflow of PAL for in situ profiling in HMC3 cells. Cells are treated with vehicle (ethanol, EtOH) or a lipid probe for 3 h. Cells are then UV irradiated, lysed, and lipid probe-labeled proteins are conjugated to biotin-azide by CuAAC reaction. The biotin labeled proteins are then enriched with streptavidin beads, trypsin digested, and the digested peptides are labeled with TMT, followed by combination, HPLC fractionation and LC-MS/MS analysis. **c**, A plot highlighting proteins selectively enriched by erucamide-diazirine probe (Eru.da) over oleamide-diazirine probe (Ole.da) and palmitamide-diazirine probe (Pal.da). X- and y-axis represent ratios of relative protein enrichment for corresponding probes (x-axis: Eru.da/Pal.da, y-axis: Eru.da/Ole.da). **d**, Confirmation of selective engagement of TMEM19 by the Eru.da performed in HEK293T cells transiently expressing TMEM19.

While it is predicted to have six transmembrane domains, the structure and function of TMEM19 remains unknown (Fig. 4a). To further investigate the function of erucamide in myeloid cells and signaling pathways mediated by erucamide through TMEM19, we conducted a bulk RNA sequencing experiment. iPSC-MPCs cells were treated with a combination of erucamide (Eru-C_18_-pSi) or not (C_18_-pSi), with (using scrambled siRNA) or without (using siTMEM19 knockdown efficiency confirmed by qPCR, Fig. 4d) TMEM19. Differential expression analysis was performed to compare the treatment combinations. When we used enrichment analyses to compare the effect of erucamide treatment in cells without or with TMEM19, the most significantly decreased pathway involves cytokine-cytokine receptor interactions (Fig. 4b and Extended Data Fig. 4a-c). In fact, 42 genes in this pathway were activated by erucamide stimulation in scrambled cells but dysregulated in siTMEM19 cells. A number of them (e.g., PDGFRB, TNFSF18, TNFSF13B and TNFRSF8) are reported to be highly associated with angiogenesis and vascular homeostasis^28–30^ (Fig. 4c). qPCR confirmed that the effective downstream cytokines/growth factors (VEGFa, PDGFα, FGF, TNFα, IGF1 and BDNF) of these pathways are up regulated by erucamide, but they were dysregulated when we knocked down TMEM19 (Fig. 4d). This aligns well with our results indicating an upregulation of pro-angiogenic genes in erucamide-injected retinas. To demonstrate if TMEM19 is essential for the activation of MPCs by erucamide stimulation, we knocked down another two top candidates from the same PAL list. It was only when TMEM19 was knocked down that we observed that most of the growth factors (e.g., FGF2, VEGFa and PDGFα) triggered by erucamide were not upregulated or activated (Extended Data Fig. 4d, e). Interestingly, the general activation of MPCs by TGFb1, LPS or DMOG was not affected by knocking down TMEM19 (Extended Data Fig. 4f). These results suggest that TMEM19 is an authentic binding protein for erucamide and is essential for pro-angiogenic and neurotrophic signaling pathways mediated by erucamide in MPCs.

**Fig. 4.**
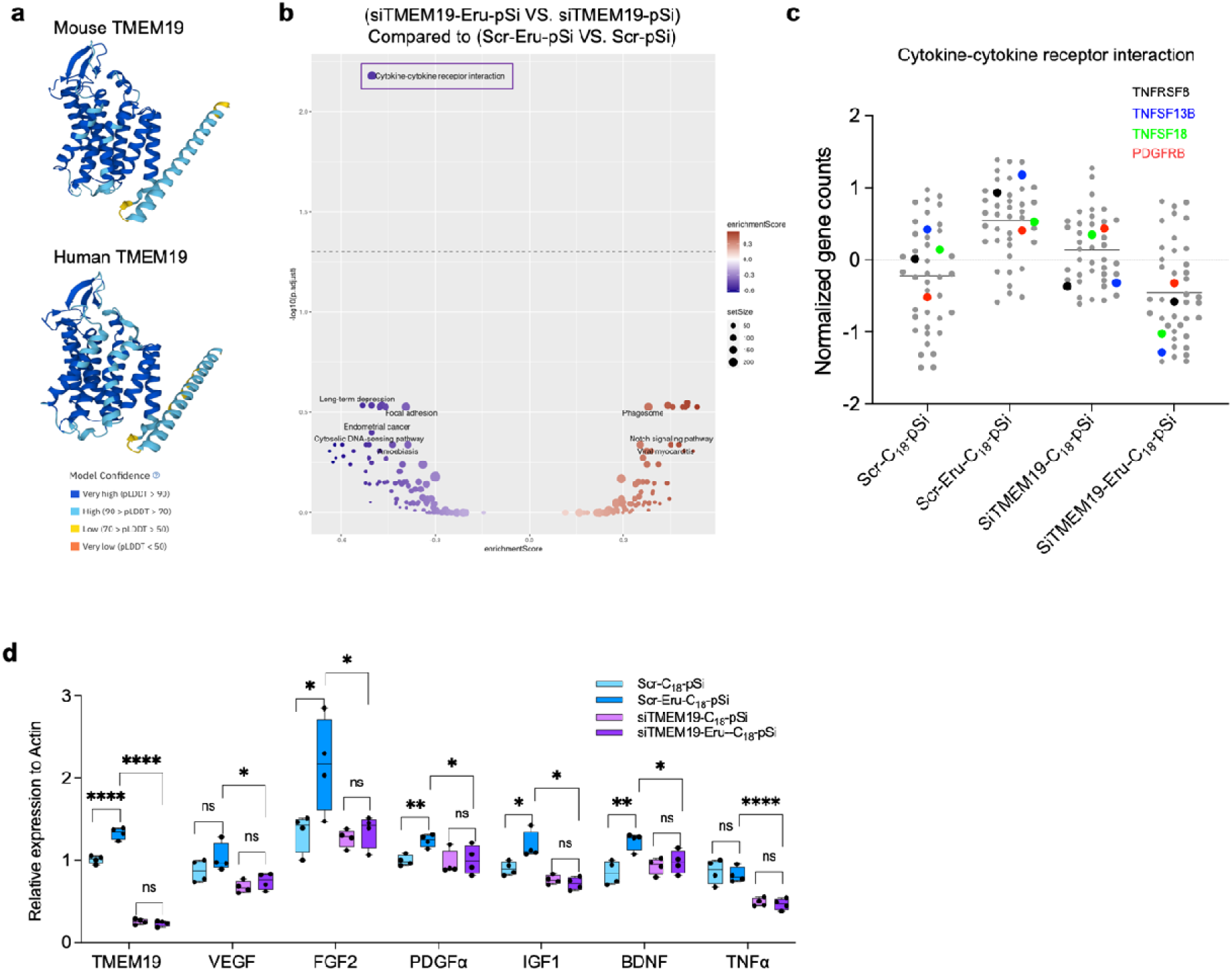
TMEM19 is a novel target protein of Erucamide in MPCs. **a**, Predicted protein structures of mouse and human TMEM19 showed 6 transmembrane domains by Alpha fold AlphaFold Protein Structure Database ^31^. **b**, KEGG Enrichment analysis of pathways changed by erucamide-TMEM19 in RNA-seq. Dysregulated cytokine-cytokine receptor pathways are labeled in purple (p. adjust=0.0066). **c**, Normalized gene counts of cytokine-cytokin receptor interaction by RNA-seq indicate changes of TNFRSF8, TNFSF13B, TNFSF18 and PDGFRB in angiogenesis related pathways. (Representative genes are labeled in colors) **d**, Results of qPCR analysis of direct effective cytokines and growth factors in MPSc used for RNA-seq (n=4).

We interfered with the pathway mediated by erucamide-TMEM19 in RD10 mice. We used neutralizing antibodies for the downstream angiogenic and neurotrophic factors of VEGFA, FGF2 and TNFα, which were found to be regulated by erucamide both in the MPCs and *in vivo* CD11b+ cells, to test if these factors are the downstream agents of erucamide rescue in degenerating retinas. The neutralizing antibodies were intravitreally injected together with Eru-C_18_-pSi as illustrated in Fig. 5a. The rescued phenotype of retinal thicknesses and deep plexus vessels induced by erucamide in RD10 mice were largely blocked by these neutralizing antibodies (Fig. 5b). Neutralizing antibodies prevented both the morphological and functional rescue by erucamide (Fig. 5 c-e), indicating the recue effects of erucamide in RD mice are mediated by erucamide-TMEM19 through the downstream angiogenic and neurotrophic factors of VEGFa, FGF2 and TNF_α_.

**Fig. 5.**
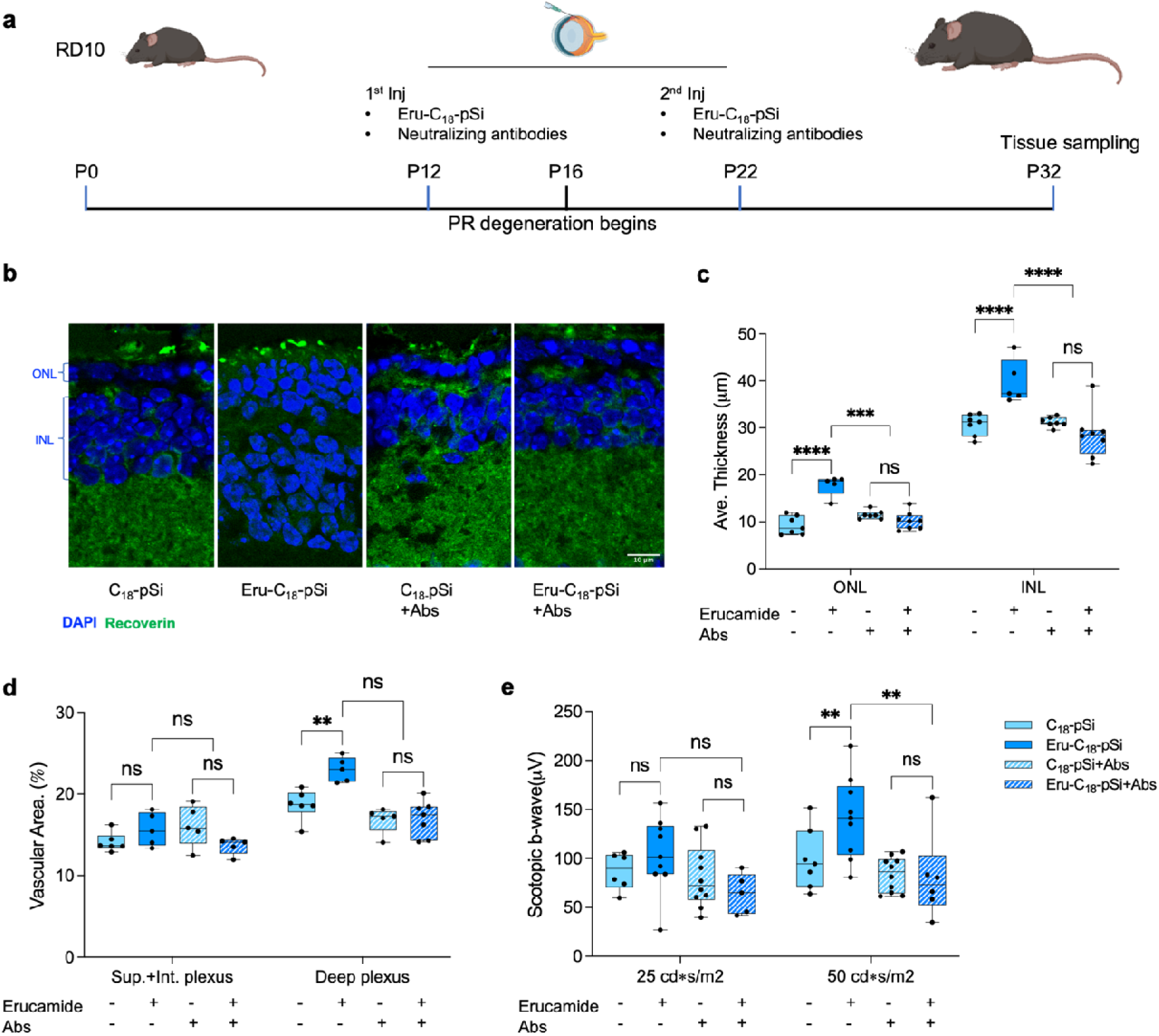
Neutralizing antibodies inhibit Erucamide’s ability to rescue retinal atrophy in RD 10 mice. **a**, Schematic illustration of Eru-C_18_-pSiNPs (214ng) together with neutralizing antibody injection in RD10 mice. **b**, Retinas from P32 RD10 mice. Photoreceptors were immunolabeled green (recoverin). **c**, Retinal thicknesses and **d**, retinal vessel density quantified in (**b**) (n>=5; experiment repeated three times.) **e**, Quantification of functional scotopic b-wave study by electroretinography.

We found that erucamide is a highly abundant in normal retinas and is severely dysregulated during retinal degeneration. Erucamide was found to be a potent pro-angiogenic, paracrin signaling molecule that regulates the retinal angiogenic environment and functions as a neurotrophic factor largely by targeting and activating MPCs through TMEM19. Knockdown of TMEM19 by siRNA results in failure to respond to the erucamide-stimulated paracrine effect of growth factors, which is consistent with *in vivo* inhibition of erucmaide-TMEM19 downstream pathways (Fig. 6). These observations provide insight into the action of the angiogenic factor erucamide as a cross-tissue signaling molecule. It may function similarly in multiple organ systems where it plays a critical role in regulating neurovasculoglial crosstalk.

**Fig. 6.**
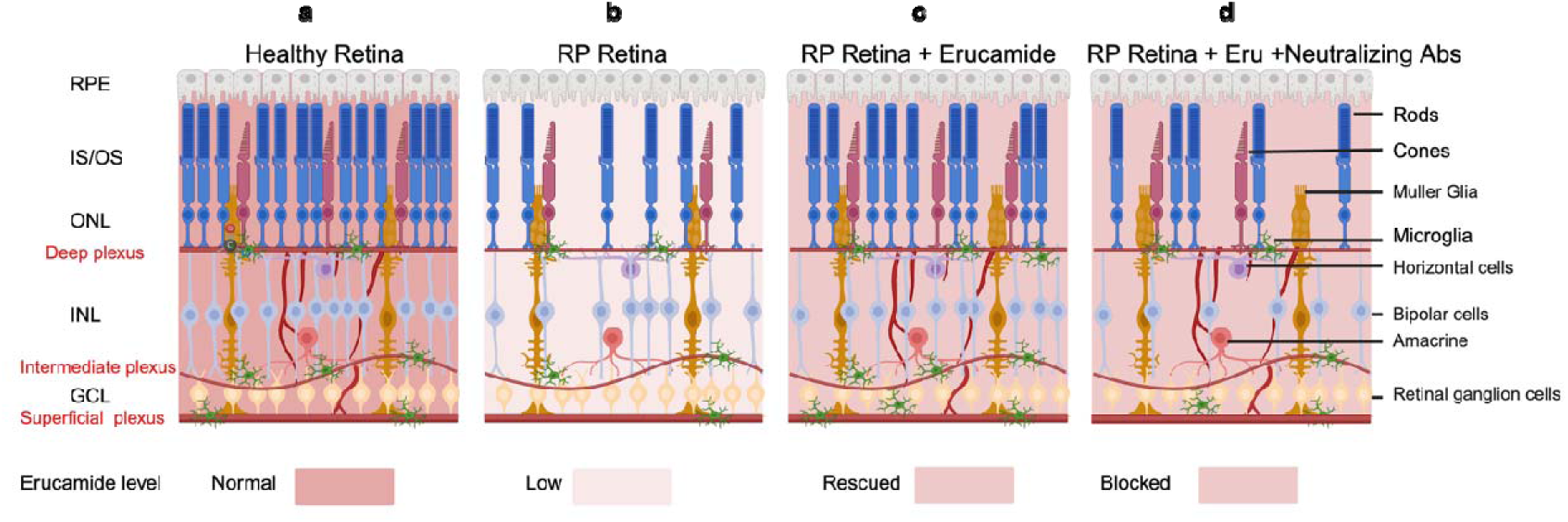
Proposed function of erucamide in the retina. **a**, In healthy retinas, a certain level of erucamide is essential to nourish the neural layers and retinal vessels by maintaining the secretion of neurotrophic and angiogenic factors from microglia. **b**, In retinas with retinitis pigmentosa, erucamide is diminished, photoreceptors are degenerated, and retinal vessels are atrophic in the deep plexus. **c**, In RP retinas rescued with erucamide injections, erucamide regulates the secretion of neurotrophic and angiogenic factors from microglia, then retinal vessel densit in the deep plexus increases and the thickness of both ONL and INL increased. **d**, In RP retinas injecte simultaneously with erucamide and downstream neutralizing antibodies, the rescue effects of erucamide are blocked. Photoreceptors are degenerate, and retinal vessels atrophy in the deep plexus as in (**b**).

## Discussion

In recent years, there has been growing recognition that neurovasculoglial crosstalk is critical to maintaining healthy, functioning neuro-vasculature in the CNS^32^, including the retina. Therapeutic approaches to preventing neuronal degeneration leading to vision loss include targeting a variety of neurovasculotrophic cytokines as well as using retinal tissue derived from induced pluripotent stem cells (iPSCs) to replace degenerated RPEs and photoreceptors to rescue degenerating retinas and visual function^33–36^. A number of studies have also demonstrated that transplanting stem cells into the eyes of retinitis pigmentosa animal models can improve visual function.^37, 38^ However, concerns persist regarding the long-term efficacy and safety of iPSC-RPE, particularly with regard to oncogenic mutations and tumorigenesis observed in preclinical models and human patients^39, 40^. For example, a pilot study placed mesenchymal stem cells, adipose stem cells, and platelet-rich plasma into the suprachoroidal space of 21 patients with RP and found only non-significant improvements in visual performance six months after treatment.

The limited survival rates and potential immune responses pose significant hurdles to the widespread clinical application of stem cell therapies in treating retinal degenerative diseases like AMD and RP. Given these limitations, we explored whether small molecules expressed by iPSC-derived retinal tissue could provide neurovasculotrophic rescue in models of retinal degeneration. Metabolomic and lipidomic screens of degenerating retinas revealed a PFAM, erucamide, with paracrine neurotrophic activity that exerted a profound rescue effect on models of retinal neurovascular degeneration. Our approach involved the use of advanced nanoparticles to allow effective *in vivo* delivery of erucamide, circumventing its hydrophobic nature and enhancing its bioavailability in the retina.

Furthermore, we identified CD11b+ myeloid cells as primary targets of erucamide within the retina. Further mechanistic studies employing photoaffinity labeling, click chemistry for biotin-derived affinity enrichment, and multiplexed quantitative proteomics alongside closely related PFAMs revealed TMEM19 as the novel transmembrane protein serving as a major and selective binding partner for erucamide. This interaction mediates downstream signaling pathways involved in neurotrophic and angiogenic factor secretion, thereby maintaining retinal homeostasis under degenerative conditions.

TMEM19 is a novel transmembrane protein with no reported function in any other cell type or tissues. Using single-cell RNA sequencing databases from the Spectacle Platform, we found that TMEM19 has low expression in retinas across different databases for humans and mice, with signals mainly from the microglia^42^, which is consistent with our discovery. Future investigations will more closely examine the precise mechanisms regulated by TMEM19 and explore additional applications of erucamide in both *in vitro* and *in vivo* models. Furthermore, the optimization of erucamide analogs to enhance solubility and prolong *in vivo* half-life could potentially offer novel small molecule-based therapies for AMD, RP, and other neurovasculoglial degenerative diseases. Conversely, the development of erucamide antagonists or TMEM19 inhibitors could provide strategies to mitigate pathological angiogenesis associated with conditions like diabetic retinopathy and neovascular AMD.

In conclusion, our findings support the use of small molecule-based therapies targeting the erucamide-TMEM19 pathway as a promising therapy for the treatment of retinal degenerative diseases, offering potential advantages in terms of safety, efficacy, and clinical applicability. This discovery expands our understanding of PFAMs beyond their established roles in the central nervous system, shedding light on their potential therapeutic applications in retinal degenerative diseases. By elucidating the molecular underpinnings of erucamide’s neurotrophic effects, this study may advance therapeutic strategies that preserve vision and improve quality of life for patients affected by these devastating conditions worldwide.

## Methods

### Animal studies

Animals were used in strict accordance with ethical guidelines of the Scripps Research Institute. RCS rats were obtained from Dr. Matthew LaVail, UCSF. RD10 mice (B6. CXB1-Pde6brd10/J) are available through the Jackson Labs. Subretinal and intraretinal injections were performed as described previously.^43, 44^ The following *in vivo* neutralizing antibodies were injected in RD10 mice (vendor, Catalog#, and concentration): TNFα (R&D systems, MAB410, 250 ng/μL), FGF-2 (05-117, Sigma-Aldrich, 1 μg/μL), and VEGF_164_ (R&D systems, AF-493-NA, 200 ng/μL).

### Ocular metabolite extraction and sample storage

Rodents were euthanized and whole eyes were dissected in PBS saline. Fresh eyes were snap frozen with dry ice and metabolites were extracted with an extraction solvent consisting of a mixture of methanol/chloroform/water/acetic acid in 60:25:14:1 (v/v), which gives a fully miscible solvent. Briefly, after the eye was transferred into a clean 2 mL glass homogenizer (Wheaton, USA), it was homogenized for 30 seconds. 200 μL of a chilled (4 °C) methanol-based extraction solvent was then added into the tissue, followed by continued homogenization for 90 seconds. Afterwards the homogenate was transferred into a tube on dry ice. Another two cycles of homogenization were repeated by adding 600 μL and 400 μL of extraction solvent, respectively. At the end of each repeat, the homogenizer was rinsed to reduce the sample loss. The whole procedure requires approximately 5 minutes, 1200+ μL of total volume was obtained at the final step (the extra μL is from the eye fluid). Then the homogenate vial was placed into liquid nitrogen for 1 min. The vial was stored on dry ice, until all tissues homogenizations are complete, homogenates were thawed on water/ice (4 °C), then centrifuged for 10-15 min at 13,000 rpm under 4 °C. The supernatant collected (sitting at -20 °C for overnight is preferable but optional) in a clean vial. The combined supernatant was dried down using a speedvac without heating. The sample is re-dissolved in ACN/Water (1:0.8) containing 5% MeOH, 5% chloroform, and 0.1% formic acid (all in volume ratio). Samples were vortexed rigorously or sonicated for 1 min before sitting it at 4 °C for ∼ 1 hour. Samples were centrifuged again, and the supernatants were transferred to LC vials for MS analysis.

### LC-MS

Analyses were performed using a 1260 HPLC system (Agilent Technologies) coupled to a 6538 UHD Accurate-Mass Q-TOF (Agilent Technologies) operated in positive (ESI+) and negative (ESI-) ionization modes. Solutions containing extracted metabolites were kept at -80°C prior to LC-MS analysis. Standards or eye extracts were separated using a Waters Xbridge C18, 3.5mm, 135A, 150m x 1.0 mm i.e. When the instrument was operated in positive ionization mode, the solvent system was A = 0.1% formic acid in water, and B = 0.1% formic acid in acetonitrile. The gradient elution starts with 100% A for injection at 0 min, increases to 80% A at 5 min and reaches 0% A at 43 min, then decreases B from 100% down to 98% from 43 to 55 min, then drops to 5% at 60 min. The injection volume was 4uL. ESI conditions included a gas temperature of 325°C, drying gas 11 L/min, nebulizer 30 psig, fragmentor 124 V, and skimmer 65 V. The instrument was set to acquire over the m/z range 25−1500, an acquisition rate of 1.3 spectra/s. MS/MS was performed in auto mode, and the instrument was set to acquire over the m/zrange 25−1500, an acquisition rate of 2 spectra/s for MS, 1.5 spectra/s for MS/MS with a iso width (the width half-maximum of the quadrupole mass bandpass used during MS/MS precursor isolation) of 1.3 m/z. MS/MS scan 5 max precursor ions at the same time, it stops after 3 scan for the same ions, then released after 0.4 min. A cycle time is 3.9s.

### LC-MS data processing and statistics

LC-MS data from the retina extract were processed using the bioinformatics analysis software XCMS Online (https://xcmsonline.scripps.edu/). The experimental MS/MS spectral data (collision energy 20 eV) was compared with the reference erucamide MS/MS spectra from the METLIN database (collision energy 20 eV)^15, 45, 46^. MS/MS spectra were manually processed by extracting ion chromatograms Qualitative Analysis of MassHunter Workstation (Agilent Technologies), pNLC and BPC extraction were performed using the MassHunter.

### Histological examinations

For histology eye cups were fixed in 4% paraformaldehyde plus 1.5% glutaraldehyde in 0.1M sodium cacodylate buffer overnight at 4°C followed by rinsing in 0.1 M Na cacodylate buffer for 1 h. Eye cups were postfixed in 1% osmium tetroxide in 0.1 M sodium cacodylate buffer for 2 h, then dehydrated in graded ethanol solutions. The tissues were incubated overnight in a 1:2 mixture of propylene oxide and Epon/Araldite (Sigma-Aldrich) and placed in 100% resin followed by embedding. The blocks were sectioned and stained with Trypan Blue. Immunohistochemistry was performed using standard protocols.^43^ The following antibodies were used (host, vendor): collagen IV (rabbit; Millipore), Recoverin (rabbit, Millipore), and CD11b (rat, BD). Secondary antibodies were all obtained from Life Technologies and DAB staining kit was purchased from Vector Labs. GS Lectin (ogies Isolectin GS-IB4 From Griffonia simplicifolia, Alexa Fluor® 568 Conjugated, I21412) was purchased from Thermofisher.

### Preparation of porous silicon (pSi)

Freestanding pSi flakes were purchased from Trutag Inc., prepared using a previously developed “perforated etch” method.^47^ The pSiNPs were obtained by immersing the pSi flakes in ethanol in a sealed vial, which was then immersed in an ultrasonication bath (Ultrasonic bath, model 97043-960, VWR International) and subjected to ultrasonic fracture conditions for 24h. The nanoparticle suspension was centrifuged at 1,500 xg for 20 min to remove large particles at the bottom of the tubes. The nanoparticle suspensions were then centrifuged at high speed (10,000 xg, 20 min) to pelletize the nanoparticles. The supernatant was discarded to eliminate the smallest nanoparticle fragments that remained in the suspension, and the pellet was resuspended in fresh ethanol. The high-speed centrifugation/resuspension procedure was carried out two more times to thoroughly wash the nanoparticles. The resulting pSiNPs displayed an average hydrodynamic size of ∼170 nm.

### Surface modification of porous silicon nanoparticles (pSiNPs)

The collected pSiNPs were washed and centrifuged (15,000 xg, 15 min) twice with toluene to remove residual ethanol. The pSiNPs were then resuspended in toluene and transferred into a glass vial, and n-octadecylsilane was added. The final concentration of pSiNPs and n-octadecylsilane was 3 mg/mL and 0.6 mg/mL respectively. A reaction vessel containing the pSiNPs and n-octadecylsilane was heated in a 95 °C oil bath for 24h with magnetic stirring. The mixture was sonicated occasionally during the reaction to ensure full suspension of the nanoparticles. The particles were subsequently washed and centrifuged three times with toluene and three times with ethanol and finally resuspended in 100% ethanol for storage.

### Characterization of pSiNPs

Hydrodynamic diameter (size) and the ζ-potential (zeta potential, or surface charge) measurements were obtained on a Zetasizer, Zs90 (Malvern Instruments). All pSiNP, C_18_-pSiNP and Eru-C_18_-pSiNP size measurements and ζ-potential measurements were measured with particles dispersed in deionized water. For hydrodynamic size measurements, 400μL of 10 µg/mL of particles were used while for ζ-potential, 800μL of 10 µg/mL of particles were used. The FTIR spectra of the particles were obtained with a Thermo Scientific Nicolet 6700 FTIR instrument fitted with a Smart iTR diamond ATR fixture. TEM images were obtained using a JEOL 1400 plus electron microscope (JEOL USA, Inc.) operating at 80KeV and subsequently imaged with a Gatan Oneview camera (Gatan, Inc.). Thermogravimetric analysis (TGA) data were obtained using a TA Instruments™ Discovery SDT 650™. Nanoparticle samples were heated from 30 °C-950 °C at a ramp rate of 10°C/min in a pure oxygen atmosphere. Both weight (%) change and derivative heat flow values were assessed, and erucamide content was determined by difference in mass loss between empty and erucamide-loaded organosilane-modified pSiNPs. Dye-loaded particle constructs were assessed using a Molecular Devices™ SpectraMax® iD5 Multi-Mode Microplate Reader. N_2_ adsorption/desorption isotherms were obtained using dry nanoparticles at a temperature of 77 K using a Micromeritics ASAP 2020 instrument.

### Loading of erucamide to pSiNPs

A 10 mg/mL stock solution of erucamide was prepared by dissolving erucamide in ethanol overnight. A 3.4 mg/mL sample of C_18_-pSiNPs in ethanol was centrifuged (15,000 xg, 10 min) to remove free ethanol and the particles were re-dispersed in 1 mL of the 10 mg/mL erucamide stock solution. The particles were then subsequently allowed to mix at 20 °C overnight. The particles were then centrifuged (15,000 xg, 10 min) and washed 3 times with PBS to remove any unbound erucamide and were resuspended in PBS or media and were used immediately after resuspension.

### Cell culture

HMC3 cell line (CRL-3304™) was purchased from ATCC and cultured with Eagle’s Minimum Essential Medium (EMEM, 30-2003™, ATCC). HEK293T cell line (CRL-3216) was purchased from ATCC and cultured with Dulbecco’s Modified Eagle’s Medium (DMEM, 30-2002 ^™^, ATCC). IPSC MPCs were derived from peripheral blood mononuclear cells from a female. The cell reprogramming was performed by the Harvard iPS core facility utilizing a sendai virus for reprogramming factor delivery. iPSCs were maintained on Cultrex (R&D systems) coated plates immersed with mTesR+ medium (STEMCELL Technologies). The cells were passaged every 3 days around 80% confluence. Colonies containing clearly visible differentiated cells were marked and manually removed prior to any subsequent passaging. All MPCs were generated using previously described methods.^25, 26^ Primary Human Umbilical Vein Endothelial Cells (HUVEC) (PCS-100-010 ™) were purchased from ATCC and cultured with Vascular Cell Basal Medium (PCS-100-030) with Endothelial Cell Growth Kit-VEGF (PCS-100-041). Human Choroidal Endothelial Cells (36052-03) were purchased from Celprogen and cultured with Human Choroidal Endothelial Cells Complete Growth Media with Serum (M36052-03S). Human Retinal Endothelial Primary Cell (36205-33) were purchased from Celprogen and cultured with Human Retinal Endothelial Cell Culture Media with Serum (M36205-33S). All cells were obtained with verified normal karyotype and contamination free.

### Photoaffinity labeling method^27^

HMC3 cells and HEK293T cells were treated with vehicle (EtOH) or a lipid probe for 3h in the incubator. HMC3 cells were then UV irradiated, lysed, and lipid probe-labeled proteins were conjugated to biotin-azide by CuAAC reaction. The biotin labeled proteins were then enriched with streptavidin beads, trypsin digested, and the digested peptides were labeled with TMT, followed by combination, HPLC fractionation and LC-MS/MS analysis. HEK293T cells were then UV irradiated then UV irradiated, lysed, and lipid probe-labeled proteins were conjugated to Rhodamine by CuAAC reaction. The labeled proteins were then used for visualising by western blot.

### Retinal myeloid cells isolation by flow cytometry

A postnatal neural dissociation kit (Miltenyi, 130-092-628) was used to prepare a single cell suspension from mouse retinas. Cells were centrifuged at 150g for 5 min at 4°C. The digested tissue was resuspended in 100 μL of 4% FBS in PBS containing an FITC antibody to CD11b (1:100; BioLegend, 101206) and incubated for 20 min on ice. The cells were washed and suspended with 1 ml of 4% FBS/PBS containing DAPI (1:2000; Thermo Fisher Scientific, 62248). Labeled retinal myeloid cells (CD11b positive) were isolated by fluorescence-activated cell sorting (FACS) (MoFlo Astrios EQ; Beckman Coulter) at the Scripps Flow Cytometry Core Facility. Sorted cells were resuspend in 350ul of RLT buffer from RNeasy Micro Kit (QIAGEN) and stored at −80°C.

### RNA isolation, Real-Time PCR and RNA-seq

For whole retina and culture cells, single retinas were collected in 500 μL of Trizol and total RNA was isolated using a PureLink RNA Mini Kit (Thermo Fisher Scientific) according to manufacturer’s instructions. Seven hundred and fifty nanograms of RNA was used for RT-qPCR using a high-capacity cDNA reverse transcription kit (Thermo Fisher Scientific). For flow-sorted cells, total RNA was isolated from sorted cells using the RNeasy Micro Kit (QIAGEN) and reverse transcribed using Maxima First Strand cDNA Synthesis Kit for RT-qPCR (Thermo Scientific). qPCR was performed using Power-up SYBR™ Green PCR Master Mix (Thermo Fisher Scientific) and primers on a Quantstudio 5 Real-Time PCR System (Thermo Fisher Scientific). β-actin (Actb) was used as the reference gene for all experiments. Levels of mRNA expression were normalized to those in controls as determined using the comparative CT (ΔΔCT) method. RNA samples were used for RNA sequencing in Scripps Genetic Cores. Quality control of raw sequencing data was performed using FastQC and MultiQC.^48^ Raw reads were aligned to the reference genome using RSubread^49^. Gene expression levels were quantified using featureCounts. Post-alignment QC included assessing the distribution of reads across genomic features (e.g., exons, introns) and checking for potential batch effects using PCA. Differential expression analysis was carried out using the limma-voom pipeline^50^. Data was filtered and normalisation was performed using the Trimmed Mean of M-values method. Linear modelling and empirical Bayes moderation were applied to identify differentially expressed genes. Genes with an FDR-adjusted p-value < 0.05 were considered significantly differentially expressed. Over-representation analysis and functional enrichment analysis were performed using the clusterProfiler^51^ package using Gene Ontology and KEGG to define biologically relevant pathways enrichment analysis was conducted.

### Confocal microscopy and quantification

All images were acquired with a confocal laser-scanning microscope (LSM 700 or 710, Carl Zeiss) and processed with the ZEN 2010 software (Carl Zeiss), ImageJ (NIH), and Photoshop CS6 (Adobe). Manual segmentation using the Adobe Photoshop CS6 ruler instrument was used to accurately measure retinal layers. Quantification of degeneration in RD10 mice (injected at P14 and analyzed at P18, P25, and P32) was also performed as follows: Eyes were cut in 14μm sections and immunolabeled with recoverin antibodies and DAPI. Retinal thickness values were measured in nine distinct locations from micrographs in central and peripheral regions in erucamide and empty microparticle injected eyes (n=4). Averages were calculated and plotted using Excel. The experiment was repeated four times. Two-tail students t-tests were used for statistical analyses.

### Ganzfeld electroretinography

Mice were dark adapted overnight, and all work was done under red light as previously described^52^.

### Probe synthesis

Methods, experimental procedures for syntheses of compound probes (**^BDP^Erucamide**, **Eru.da**, **Ole.da** and **Pal.da**) are listed in **Supplementary information**.

### Statistical analysis

Statistical analyses were performed using GraphPad Prism 9.10 software. Unless indicated otherwise, the evaluated parameters in the groups are shown as the means and standard deviations (s.d.) or standard error of the mean (s.e.m.) for each group. All the results were compared using two-sided independent t-test to compare two groups, one-way ANOVA with Fisher’s least significant difference post hoc test to compare more than two groups, and two-way ANOVA with Fisher’s least significant difference post hoc test to compare two-factor study design. For all tests, *p*< 0.05 was considered significant. All data points in the manuscript represent individual biological replicates. No statistical methods were used to predetermine sample size.

## Code Availability

This paper does not report the original code.

## Acknowledgements

We thank the Lowy Medical Research Institute for funding this work. The initial phases of this work were also funded by grants from the National Eye Institute (R01EY11254 and 5R24EY017540) and the California Institute for Regenerative Medicine (TR1-01219) to MF. Additional funding came from the NSF through the UC San Diego Materials Research Science and Engineering Center (UCSD MRSEC), grant DMR-2011924, by the National Institutes of Health, through grant 2R01AI132413. This work was also performed in part at the San Diego Nanotechnology Infrastructure (SDNI) of UCSD, a member of the National Nanotechnology Coordinated Infrastructure, which is supported by the National Science Foundation (Grant ECCS-2025752).

The authors acknowledge the use of facilities and instrumentation supported by NSF through the UC San Diego Materials Research Science and Engineering Center (UCSD MRSEC) DMR-2011924. S.V. acknowledges financial support from the Natural Sciences and Engineering Research Council of Canada Postgraduate Scholarship—Doctoral program (NSERC PGS-D).

We also thank Lea Scheppke, Michael Dorrell and Marin Gantner for editing this paper.

## Author Contributions

G.S., B.C., P.W., S.V., M.J.S., D.B., and M.F. conceived and designed the study. Data acquisition and processing was performed by G.W., S.C., Q.Y., S.V., D.O., S.G., P.W., J.W., R.F., H.P., E.A., J.R., A.O., R.B., and K.E. Data were visualized G.W., S.C., Q.Y., D.O., P.W., J.W., and R.B. Data interpretation was done by G.W. Funding acquisition was done by M.J.S. and M.F. Writing of the original draft was performed by G.W., S.C., K.E., and M.F. All authors contributed to the review and editing of this manuscript.

## Materials & Correspondence

Any material requests and additional information needed to reanalyze the data reported in this paper is available from the corresponding author— Martin Friedlander upon reasonable request.

---

**Extended Data Fig. 1.**
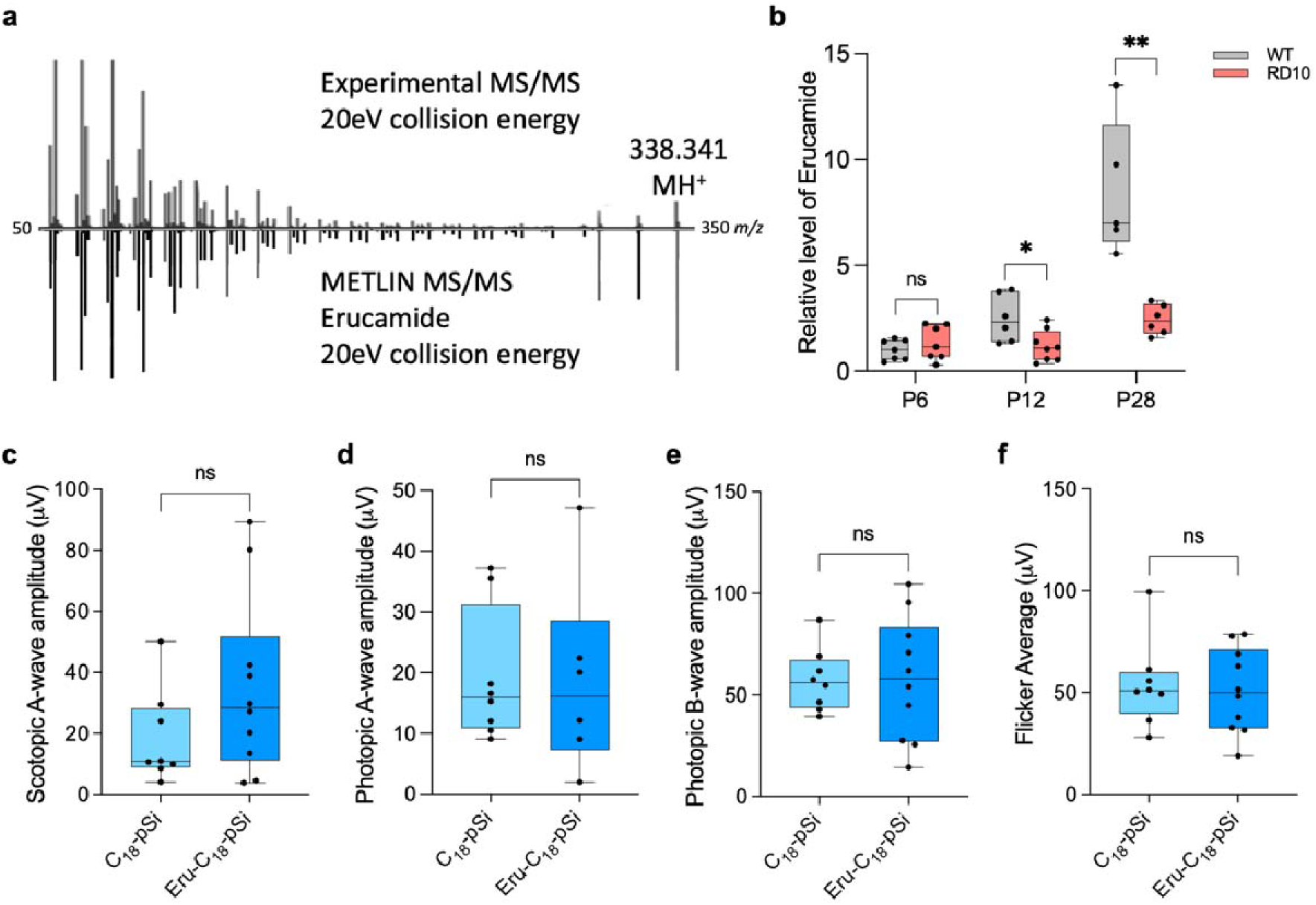
Erucamide decreased with retinal degeneration. **a**, The experimental MS/MS spectral data (collision energy 20 eV) was compared with the reference erucamide MS/MS spectra from the METLIN database (collision energy 20 eV)^15, 45^. The comparison revealed highly consistent fragmentation patterns in both the intensity ratios and accurate masses of the fragment ions. **b**, Erucamide level is attenuated in RD10 at P12 (n>=6; p=0.021). and P28 (n>=5; p=0.0013). **c**, Electroretinographic (ERG) measurements of scotopic A-wave responses to a flash in dark-adapted RD mice injected with Eru or vehicle at P32 (n>=8 per group). **d**,**e**,**f**, ERG measurements of photopic A-wave responses (**d**), photopic B-wave responses (**e**), and photopic flicker responses (**f**) to a flash in RD mice injected with Eru or vehicle at P32 (n>=6 per group).

**Extended Data Fig. 2.**
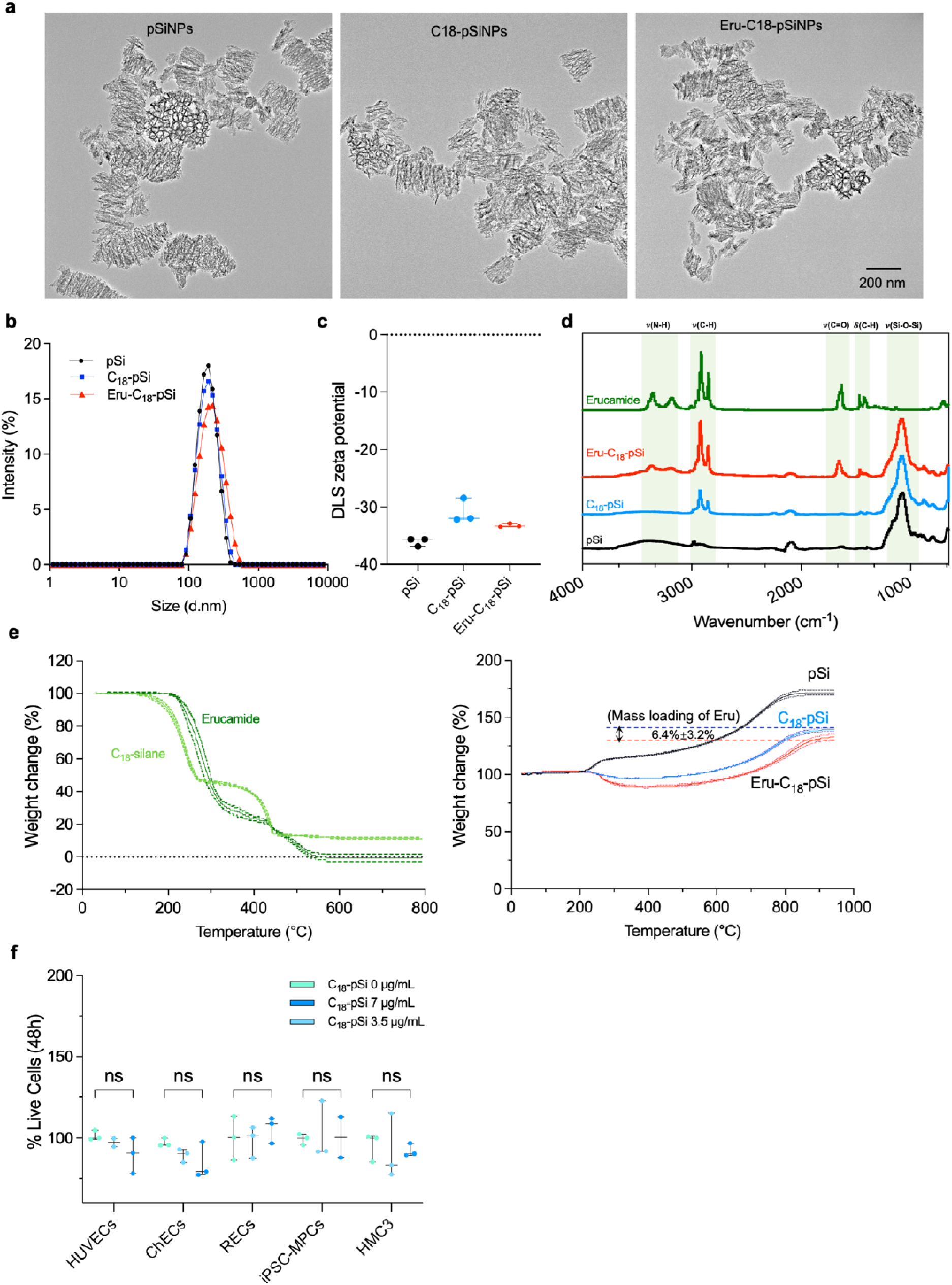
Characterization of Erucamide-loaded, silane-functionalized nanoparticles (Eru-C_18_-pSiNPs). **a**, Transmission Electron Microscopy (TEM) images of freshly etched (pSiNP) and octadecylsilane-functionalized (C_18_-pSiNP) and erucamide-loaded, octadecylsilane-functionalized particles (Eru-C_18_-pSiNP) (scale bar = 200 nm). **b**, Dynamic light scattering (DLS) data of size distribution and **c**, ζ-potential measurements. **d**, Fourier Transform Infrared Spectroscopy (FTIR) spectra of pSiNPs functionalized with silanes and loaded with erucamide. **e**, Thermogravimetric analysis (TGA) measurements of Eru-C_18_-pSiNPs. The derivative heat flow indicates phase changes of erucamide and the octadecylsilane reagent while the weight (%) curve provides a estimate of the amount of functionalized silane and loaded erucamide is available in NPs. The measurements indicated a weight percent loss of 6.4% (wt.%) of erucamide within the particles at the end of 800°C in resulting in the vaporization of erucamide. **f**, Erucamide-C_18_-pSiNPs have no toxicity in multiple cells. *In vitro* toxicity of pSiNP constructs. Viability of human umbilical vein endothelial cells (HUVECs), human choroidal endothelial cells (ChECs), human retinal endothelial cells (RECs), human iPSC-derived macrophage precursor cells (MPCs), and human microglia cell line (HMC3) were incubated with various concentrations of C_18_-pSiNP constructs for 48 hours. The cell viability was assessed using Invitrogen™LIVE/DEAD™ Viability/Cytotoxicity Kit following the manufacturer’s protocol. (n=3)

**Extended Data Fig. 3.**
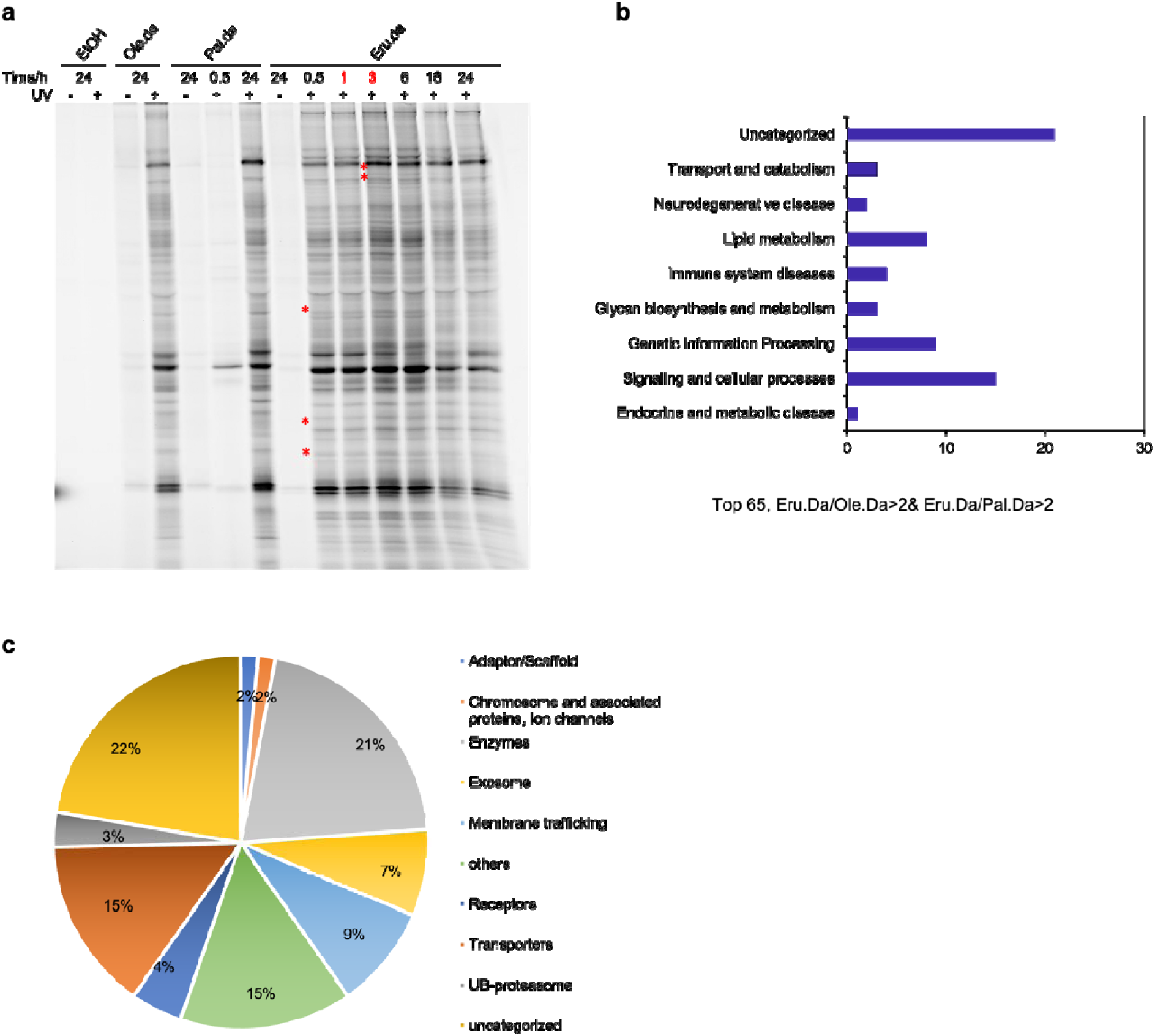
Identifying the binding protein of Erucamide in microglia by PAL. **a**, Erucamide, oleamide and palmitamide probes show overlapping but distinct protein interaction profiles in HMC3 cells (red stars indicated Eru.da specific labelled proteins). **b**, Involvement in specific biological processes of erucamide targetin proteins. **c**, Protein class distribution of erucamide targeting proteins. (**b-c**, for top 65 list: Eru.da/Ole.da>2 & Eru.da/Pal.da>2)

**Extended Data Fig. 4.**
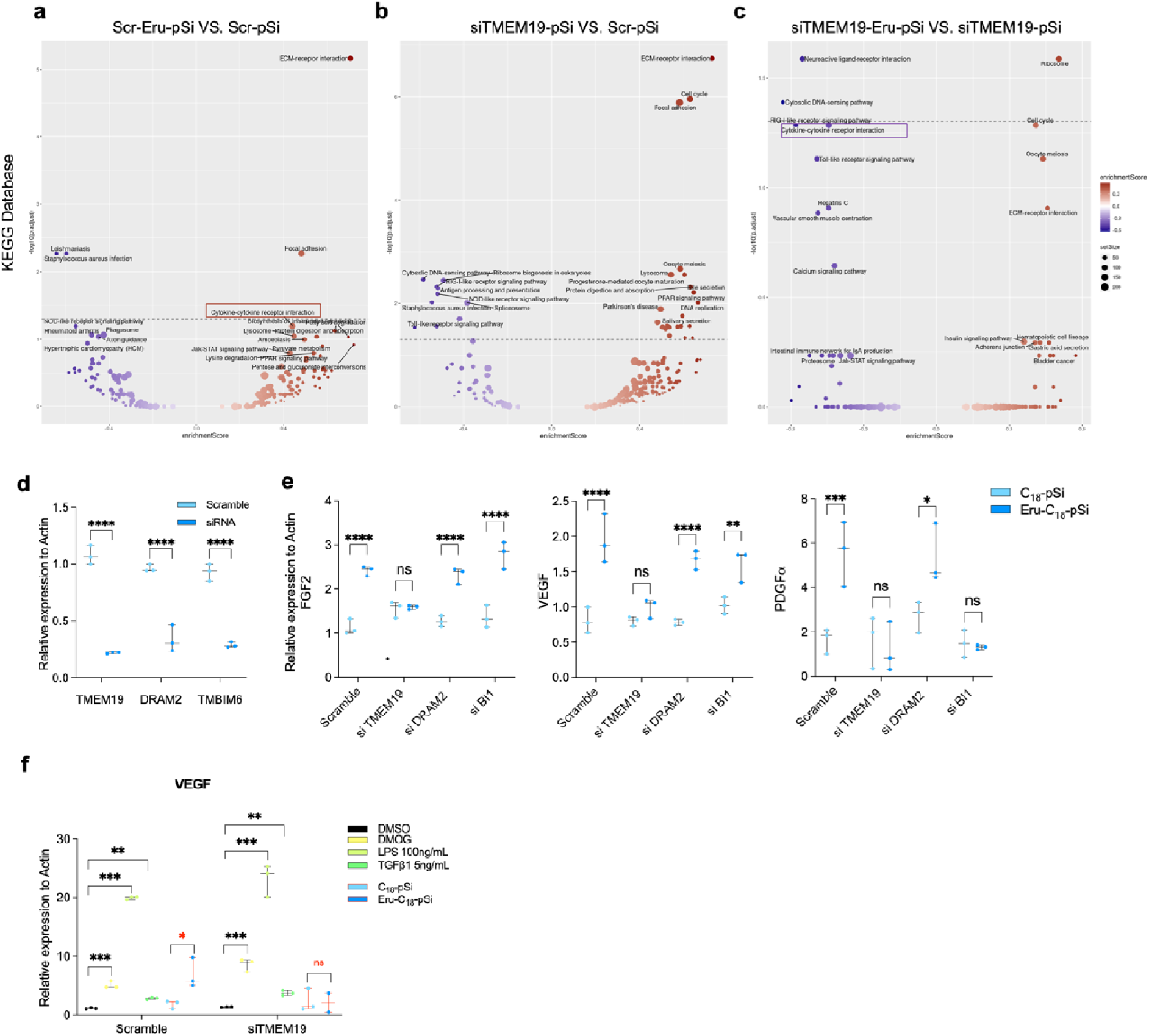
RNA-seq pathway enrichment analysis of Erucamide-TMEM19 pathway. **a**, Enrichment analysis of erucamide-mediated pathways in Scrambled MPCs compared to empty pSiNPs treatment using KEGG-database, all genes are used, the direction of effect is tested to see if pathway is upregulated or downregulated. Upregulated cytokine-cytokine receptor pathway is labeled in red box. **b**, KEGG analysis of general pathways in siTMEM19 MPCs compared to Scr MPCs. **c**, KEGG analysis of erucamide mediate pathways in siTMEM19 MPCs compared to empty pSiNPs treatment. Downregulated cytokine-cytokine receptor pathway is labeled in purple box. **d**, TMEM19 and two other candidates (DRAM2 and TMBIM6) from chem-proteomics are knocked down in iPSC-MPCs by siRNA (n=3). **e**, Angiogenic factors are not upregulated in TMEM19-deprived cells, but not in other cell groups. (n=3). **f**, The general activation of MPCs by DMOG, LPS and TGFβ1 are not affected by TMEM19 knock-down while the activation by erucamide (0.64 µM) is through TMEM19 (n=3).

**Extended Data Video. 1.**
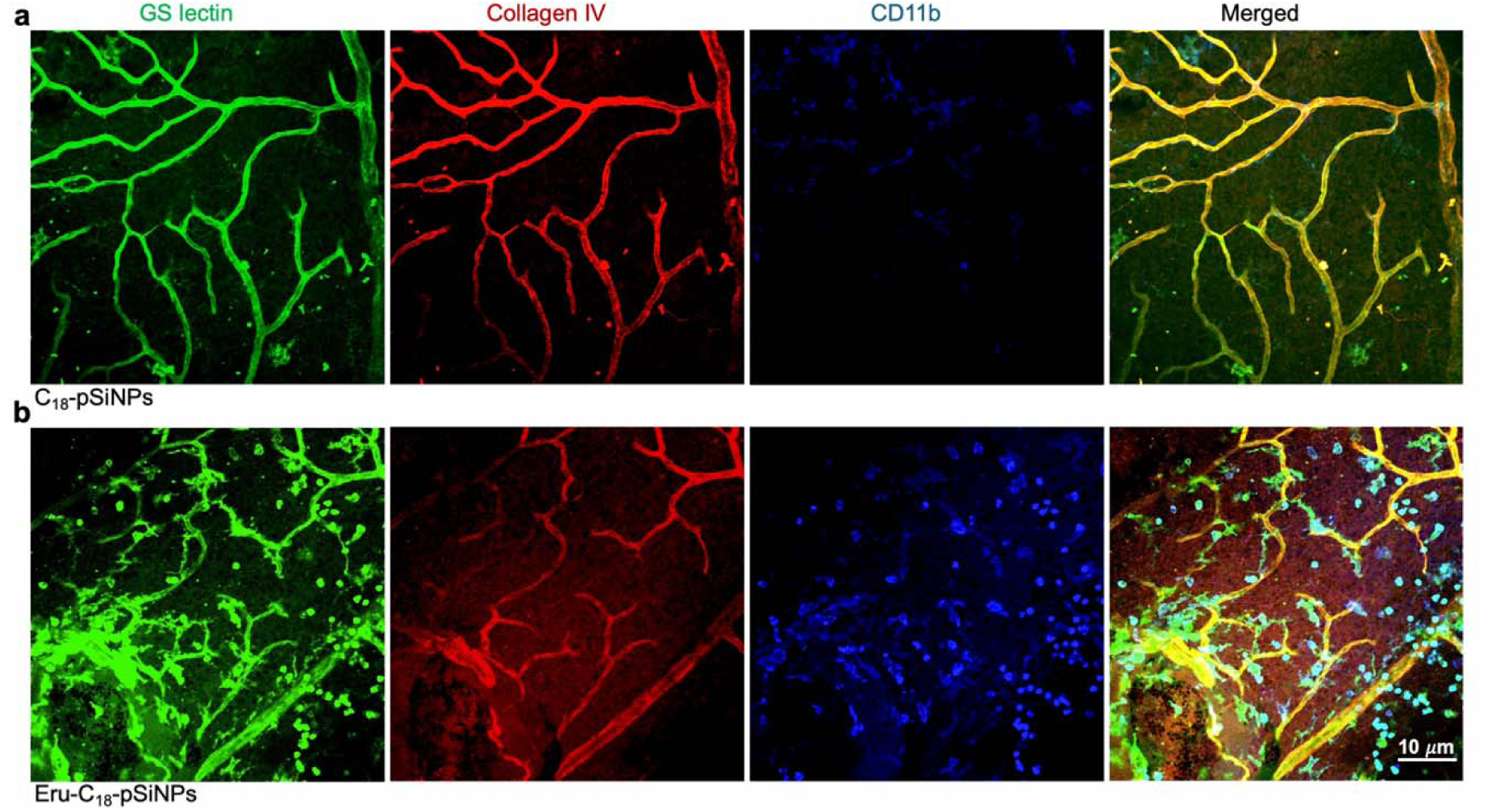
Videos of erucamide activating CD11b+ macrophage/microglia in subretinal injected retinas. **a**, C_18_-pSiNPs injected retinas exhibited no activated morphological changes in CD11b+ cells. **b**, Erucamide-C_18_-pSiNPs (214ng) injected retinas exhibited activated CD11b+ cells in all layers of superficial, intermediate, and deep plexuses.

## References

1. Verbakel, S.K., et al. Non-syndromic retinitis pigmentosa. Prog Retin Eye Res 66, 157–186 (2018).

2. Milam, A.H., Li, Z.Y. & Fariss, R.N. Histopathology of the human retina in retinitis pigmentosa. Prog Retin Eye Res 17, 175–205 (1998).

3. O’Neal, T.B. & Luther, E.E. Retinitis Pigmentosa. in StatPearls (Treasure Island (FL), 2024).

4. Strauss, O. The retinal pigment epithelium in visual function. Physiol Rev 85, 845–881 (2005).

5. Lewandowski, D., et al. Dynamic lipid turnover in photoreceptors and retinal pigment epithelium throughout life. Prog Retin Eye Res 89, 101037 (2022).

6. Gantner, M.L., et al. Serine and Lipid Metabolism in Macular Disease and Peripheral Neuropathy. N Engl J Med 381, 1422–1433 (2019).

7. Schwitzer, T., Schwan, R., Angioi-Duprez, K., Giersch, A. & Laprevote, V. The Endocannabinoid System in the Retina: From Physiology to Practical and Therapeutic Applications. Neural Plast 2016, 2916732 (2016).

8. Cravatt, B.F., et al. Chemical characterization of a family of brain lipids that induce sleep. Science 268, 1506–1509 (1995).

9. Ezzili, C., Otrubova, K. & Boger, D.L. Fatty acid amide signaling molecules. Bioorg Med Chem Lett 20, 5959–5968 (2010).

10. Arafat, E.S., Trimble, J.W., Andersen, R.N., Dass, C. & Desiderio, D.M. Identification of fatty acid amides in human plasma. Life Sci 45, 1679–1687 (1989).

11. Waluk, D.P., et al. Chapter 9 - Mammalian Fatty Acid Amides of the Brain and CNS. in Omega-3 Fatty Acids in Brain and Neurological Health (ed. R.R. Watson & F. De Meester) 87–107 (Academic Press, Boston, 2014).

12. Wakamatsu, K., Masaki, T., Itoh, F., Kondo, K. & Sudo, K. Isolation of fatty acid amide as an angiogenic principle from bovine mesentery. Biochem Biophys Res Commun 168, 423–429 (1990).

13. Mitchell, C.A., et al. Enhancement of neovascularization in regenerating skeletal muscle by the sustained release of erucamide from a polymer matrix. J Biomater Appl 10, 230–249 (1996).

14. Smith, C.A., Want, E.J., O’Maille, G., Abagyan, R. & Siuzdak, G. XCMS:[l] Processing Mass Spectrometry Data for Metabolite Profiling Using Nonlinear Peak Alignment, Matching, and Identification. Analytical Chemistry 78, 779–787 (2006).

15. Xue, J., Guijas, C., Benton, H.P., Warth, B. & Siuzdak, G. METLIN MS2 molecular standards database: a broad chemical and biological resource. Nature Methods 17, 953–954 (2020).

16. Hamberger, A. & Stenhagen, G. Erucamide as a modulator of water balance: new function of a fatty acid amide. Neurochem Res 28, 177–185 (2003).

17. Sailor, M.J. & ProQuest. Porous silicon in practice : preparation, characterization and applications (Wiley-VCH, Weinheim, 2012).

18. Kim, B., et al. Immunogene therapy with fusogenic nanoparticles modulates macrophage response to Staphylococcus aureus. Nat Commun 9, 1969 (2018).

19. Nan, K., et al. Porous silicon oxide-PLGA composite microspheres for sustained ocular delivery of daunorubicin. Acta Biomater 10, 3505–3512 (2014).

20. Kim, D., et al. Thermally Induced Silane Dehydrocoupling on Silicon Nanostructures. Angewandte Chemie 128, 6533–6537 (2016).

21. Dulal, N., et al. Slip-additive migration, surface morphology, and performance on injection moulded high-density polyethylene closures. J Colloid Interface Sci 505, 537–545 (2017).

22. Hernández-Fernández, J., Puello-Polo, E. & López-Martínez, J. Recovery of (Z)-13-Docosenamide from Industrial Wastewater and Its Application in the Production of Virgin Polypropylene to Improve the Coefficient of Friction in Film Type Applications. Sustainability 15, 1247 (2023).

23. Chang, B., et al. Two mouse retinal degenerations caused by missense mutations in the beta-subunit of rod cGMP phosphodiesterase gene. Vision Res 47, 624–633 (2007).

24. Friedlander, M., et al. Involvement of integrins alpha v beta 3 and alpha v beta 5 in ocular neovascular diseases. Proc Natl Acad Sci U S A 93, 9764–9769 (1996).

25. Usui-Ouchi, A., et al. Deletion of Tgfbeta signal in activated microglia prolongs hypoxia-induced retinal neovascularization enhancing Igf1 expression and retinal leukostasis. Glia 70, 1762–1776 (2022).

26. Usui-Ouchi, A., et al. Integrating human iPSC-derived macrophage progenitors into retinal organoids to generate a mature retinal microglial niche. Glia 71, 2372–2382 (2023).

27. Niphakis, M.J., et al. A Global Map of Lipid-Binding Proteins and Their Ligandability in Cells. Cell 161, 1668–1680 (2015).

28. Yin, L., et al. HOOK1 Inhibits the Progression of Renal Cell Carcinoma via TGF-beta and TNFSF13B/VEGF-A Axis. Adv Sci (Weinh) 10, e2206955 (2023).

29. Aggarwal, B.B., Gupta, S.C. & Kim, J.H. Historical perspectives on tumor necrosis factor and its superfamily: 25 years later, a golden journey. Blood 119, 651–665 (2012).

30. Andrae, J., Gallini, R. & Betsholtz, C. Role of platelet-derived growth factors in physiology and medicine. Genes Dev 22, 1276–1312 (2008).

31. Jumper, J., et al. Highly accurate protein structure prediction with AlphaFold. Nature 596, 583–589 (2021).

32. Alkayed, N.J. & Cipolla, M.J. The Ever-Evolving Concept of the Neurovascular Unit. Stroke 54, 2178–2180 (2023).

33. D’Antonio-Chronowska, A., D’Antonio, M. & Frazer, K.A. In vitro Differentiation of Human iPSC-derived Retinal Pigment Epithelium Cells (iPSC-RPE). Bio Protoc 9, e3469 (2019).

34. Hazim, R.A., et al. Differentiation of RPE cells from integration-free iPS cells and their cell biological characterization. Stem Cell Res Ther 8, 217 (2017).

35. Krohne, T.U., et al. Generation of retinal pigment epithelial cells from small molecules and OCT4 reprogrammed human induced pluripotent stem cells. Stem Cells Transl Med 1, 96–109 (2012).

36. Westenskow, P.D., Kurihara, T. & Friedlander, M. Utilizing stem cell-derived RPE cells as a therapeutic intervention for age-related macular degeneration. Adv Exp Med Biol 801, 323–329 (2014).

37. He, Y., et al. Recent advances of stem cell therapy for retinitis pigmentosa. Int J Mol Sci 15, 14456–14474 (2014).

38. Surendran, H., et al. Transplantation of retinal pigment epithelium and photoreceptors generated concomitantly via small molecule-mediated differentiation rescues visual function in rodent models of retinal degeneration. Stem Cell Res Ther 12, 70 (2021).

39. Aoi, T., et al. Generation of pluripotent stem cells from adult mouse liver and stomach cells. Science 321, 699–702 (2008).

40. Okita, K., Ichisaka, T. & Yamanaka, S. Generation of germline-competent induced pluripotent stem cells. Nature 448, 313–317 (2007).

41. Limoli, P.G., Vingolo, E.M., Limoli, C. & Nebbioso, M. Stem Cell Surgery and Growth Factors in Retinitis Pigmentosa Patients: Pilot Study after Literature Review. Biomedicines 7 (2019).

42. Voigt, A.P., et al. Spectacle: An interactive resource for ocular single-cell RNA sequencing data analysis. Exp Eye Res 200, 108204 (2020).

43. Westenskow, P.D., et al. Ras pathway inhibition prevents neovascularization by repressing endothelial cell sprouting. J Clin Invest 123, 4900–4908 (2013).

44. Westenskow, P.D., et al. Performing subretinal injections in rodents to deliver retinal pigment epithelium cells in suspension. J Vis Exp, 52247 (2015).

45. Guijas, C., et al. METLIN: A Technology Platform for Identifying Knowns and Unknowns. Analytical Chemistry 90, 3156–3164 (2018).

46. Zhu, Z.-J., et al. Liquid chromatography quadrupole time-of-flight mass spectrometry characterization of metabolites guided by the METLIN database. Nature Protocols 8, 451–460 (2013).

47. Kang, J., et al. Self-Sealing Porous Silicon-Calcium Silicate Core-Shell Nanoparticles for Targeted siRNA Delivery to the Injured Brain. Adv Mater 28, 7962–7969 (2016).

48. Ewels, P., Magnusson, M., Lundin, S. & Kaller, M. MultiQC: summarize analysis results for multiple tools and samples in a single report. Bioinformatics 32, 3047–3048 (2016).

49. Liao, Y., Smyth, G.K. & Shi, W. The R package Rsubread is easier, faster, cheaper and better for alignment and quantification of RNA sequencing reads. Nucleic Acids Res 47, e47 (2019).

50. Law, C.W., et al. RNA-seq analysis is easy as 1-2-3 with limma, Glimma and edgeR. F1000Res 5 (2016).

51. Wu, T., et al. clusterProfiler 4.0: A universal enrichment tool for interpreting omics data. Innovation (Camb) 2, 100141 (2021).

52. Lim, E.W., et al. Serine and glycine physiology reversibly modulate retinal and peripheral nerve function. Cell Metab 36, 2315–2328 e2316 (2024).

