## Supplementary Information for "Erucamide regulates retinal neurovascular crosstalk"

Author Information

^1^Department of Molecular Medicine, Scripps Research 10550 N. Torrey Pines Road, La Jolla, California 92037, United States

^2^Department of Chemistry, Scripps Research 10550 N. Torrey Pines Road, La Jolla, California 92037, United States

^3^Lowy Medical Research Institute, La Jolla, California 92037, United States

^4^Department of Chemistry and Biochemistry, University of California, San Diego, La Jolla, California 92093, United States

^5^Materials Science and Engineering Program, University of California, San Diego, La Jolla, California 92093, United States

^6^Department of Integrative Structural and Computational Biology, The Scripps Research Institute, La Jolla, CA 92037, USA

**Table of Contents**

Methods, Experimental procedures for syntheses of compound probes	Page

General Procedures	S1

^BDP^Eru probe	S2 – S4

Eru.da probe	S5 – S6

Ole.da probe	S7

Pal.da probe	S7

The gating strategy for FAC sorted CD11b+ cells in Fig. 2e	S8

**Experimental Procedures**

**General Procedure.** All commercial reagents were used without further purification unless otherwise noted. THF was distilled prior to use. All reactions were performed in oven-dried (200 °C) glassware and under an inert atmosphere of anhydrous Ar (argon) unless otherwise noted. Column chromatography was performed with silica gel 60. TLC was performed on Whatman silica gel (250 μm) F_254_ glass plates and compounds were visualized by UV or common stains. PTLC was performed on Whatman silica gel (250 and 500 μm) F_254_ glass plates. ^1^H NMR spectra were recorded on a Bruker 500 or 400 MHz spectrometer. Chemical shifts are measured and reported relative to an internal standard of residual CHCl_3_ (δ 7.26 for ^1^H). H^1^ NMR data are reported as follows: chemical shift (δ), multiplicity (ovlp=overlapping, br=broad, s=singlet, d=doublet, t=triplet, q=quartet, m=multiplet), coupling constant (*J*), and integration. High resolution mass spectra were obtained on an Agilent ESI-TOF/MS using Agilent ESI-L low concentration tuning mix as internal high resolution calibration standards.

**^BDP^Eru**

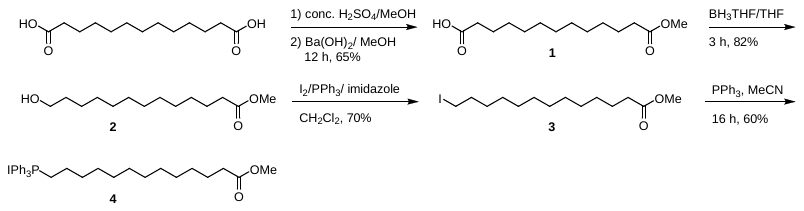

**13-Methoxy-13-oxotridecanoic Acid (1)**: Concentrated sulfuric acid (8 mL) was added to solution of tridecanedioic acid (2.00 g, 8.19 mmol) in methanol. The mixture was warmed at reflux for 2 h, cooled to room temperature and diluted with water (25 mL). The resulting mixture was exhaustively extracted with toluene. The resulting organic layer was washed with water and sodium carbonate (5% aqueous solution). The toluene layer was then concentrated to dryness under reduced pressure to yield dimethyl tridecanedioate which was used for the next step without purification.

A solution of barium hydroxide (0.5 M in anhydrous methanol, 7 mL) was added to a solution of the diester (1.73 g, 6.35 mmol) in anhydrous methanol (7 mL) in a sealed tube. The solution was vigorously stirred at room temperature for 18 h. The barium salt was then filtered, washed with methanol and dissolved in aqueous 4 N HCl. The resulting solution was extracted with diethyl ether. The organic layers were combined, and solvents were evaporated under reduced pressure. The resulting solution was purified by flash chromatography (SiO_2,_ 90% EtOAc-hexanes) to yield **1** (1.07 g, 65%) as a white solid. ^1^H NMR (CDCl_3_, 400 MHz) δ: 3.68 (s, 3H), 2.35 (t, *J* = 7.4 Hz, 2H), 2.31 (t, *J* = 2.5 Hz, 2H), 1.62 (m, 4H), 1.27 (m, 15H). LRMS-ESI *m/z* calculated for C_14_H_27_O_3_^+^ [M+H^+^] calculated: 259.91, found 259.91.

**Methyl 13-Hydroxytridecanoate (2)**: A solution of **1** (50 mg, 0.19 mmol, 1 equiv) in THF (1 mL) at 0 °C was treated with 1 M BH_3_^.^THF (0.57 mL, 0.57 mmol, 3 equiv) and stirred for 5 minutes. The reaction was quenched by the addition of NaHCO_3_ at 0 °C and slowly allowed to warm to room temperature. The resulting solution was extracted with EtOAc, dried over Na_2_SO_4_, filtered and concentrated under reduced pressure. Column chromatography (SiO_2_, 50% EtOAc-hexanes) yielded **2** as a white solid (37 mg, 82%). ^1^H NMR (CDCl_3_, 400 MHz) δ: 3.66 (s, 3H), 3.64 (t, *J* = 6.6 Hz, 2H), 2.30 (t, *J* = 7.5 Hz, 2H), 1.62 (m, 2H). 1.26 (m, 19H). LRMS-ESI *m/z* calculated for C_14_H_29_O_3_^+^ [M+H^+^] calculated: 245.21, found 245.21.

**Methyl 13-Iodotridecanoate (3)**: A solution of **2** (415 mg, 1.7 mmol, 1 equiv) in CH_2_Cl_2_ (15 mL) was treated with triphenylphosphine (624.2 mg, 2.38 mmol, 1.4 equiv) and imidazole (221 mg, 3.23 mmol, 1.9 mmol) and cooled to 0 °C. I_2_ (604 mg, 2.38 mmol, 1.4 equiv) was added in batches over 10 min and the solution was slowly warmed to room temperature. The reaction mixture was quenched with the addition of Na_2_S_2_O_3_ and extracted with CH_2_Cl_2_. The resulting mixture was dried over Na_2_SO_4_, filtered and concentrated. Column chromatography (SiO_2_, 10% EtOAc-hexanes) yielded **3** as a white solid (420 mg, 70%). ^1^H NMR (CDCl_3_, 400 MHz) δ: 3.66 (s, 3H), 3.19 (t, *J* = 7.1 Hz, 2H), 2.30 (t, *J* = 7.4 Hz, 2H), 1.26 (m, 21H). LRMS-ESI *m/z* calculated for C_14_H_28_IO_2_^+^ [M+H^+^] calculated: 355.11, found 355.11.

**((12-Methoxycarbonyl)undecyl)triphenylphosphonium Iodide (4)**: An acetonitrile (20 mL) solution of iodide **3** (1.0 g, 3.36 mmol) and triphenylphosphine (967 mg, 3.69 mmol) was warmed at reflux under nitrogen for 36 h. After cooling, the acetonitrile was removed, and the product was dissolved in CH_2_Cl_2_ (20 mL) and precipitated by dilution with ether (100 mL) to obtain 1.9 g (97%) of **4** as a white solid. ^1^H NMR (CDC1_3_, 400 MHz) δ: 7.88-7.78 **(**m, 9H), 7.77-7.69 (m, 6H), 3.70-3.45 **(**m**,** 2H), 3.65 (s, 3H), 2.26 (t, *J* = 7 Hz, 2H), 1.70-1.50 (m, 10H), 1.35-1.20 (m, 14H).

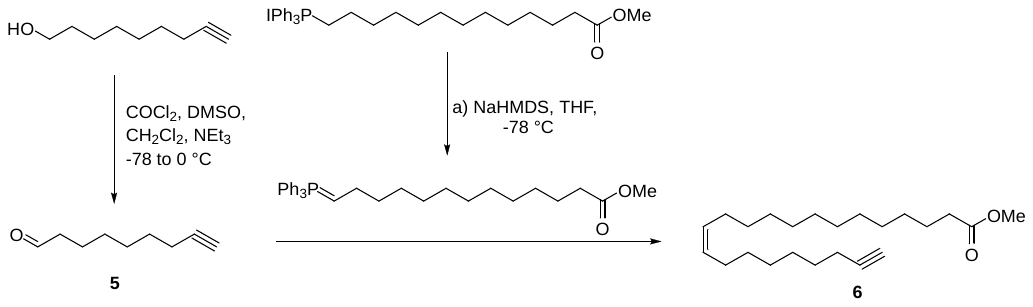

**Non-8-ynal (5)**: In an oven-dried round-bottomed flask, oxalyl chloride (0.2 mL, 2.59 mmol, 4.0 equiv) was added to anhydrous CH_2_Cl_2_ (50 mL) at –78 °C under N_2_. DMSO (0.37 mL, 5.19 mmol, 8.0 equiv) was added dropwise. After 30 min of stirring at –78 °C, non-8-yn-1-ol (100 mg, 0.65 mmol, 1.0 equiv) in anhydrous CH_2_Cl_2_ (10 mL) was added over 15 min and the reaction mixture was stirred for another 30 min. Et_3_N (0.72 mL, 5.18 mmol, 8.0 equiv) was added and the reaction mixture was stirred for 30 min at –78 °C before warming to 0 °C in an ice bath. After 30 min, the reaction was diluted with EtOAc and the reaction mixture was washed with water, dried over Na_2_SO_4_, filtered and concentrated. Column chromatography (SiO_2_, 10% EtOAc-hexanes) yielded aldehyde **5** as a yellow liquid. ^1^H NMR (CDC1_3_, 400 MHz) δ: 9.76 (t, *J* = 1.1 Hz, 1H), 2.42 (dt, *J* = 7.3, 1.1 Hz, 2H), 2.18 (dt, *J* = 6.9, 2.1 Hz, 2H), 1.63 (m, 2H), 1.52 (m, 2H), 1.33 (m, 4H). LRMS-ESI *m/z* calculated for C_9_H_15_O^+^ [M+H^+^] calculated: 139.11, found 139.11.

**Methyl (*Z*)-Docos-13-en-21-ynoate (6)**: A stirred solution of phosphonium salt **4** (208 mg, 0.34 mmol, 1.0 equiv) in anhydrous THF (15 mL) at –78 °C under N_2_ was treated with NaHMDS (0.47 mL, 0.47 mmol, 1.0 M in THF, 1.0 equiv) dropwise over 15 min. The resulting orange solution was stirred at –78 °C for an additional 1 h. In a separate round-bottomed flask, aldehyde **5** (77 mg, 0.56 mmol, 1.2 equiv) was dissolved in anhydrous THF (15 mL) and cooled to –78 °C under N_2_. The ylide solution was then slowly added to the aldehyde solution over 45 min and the mixture was stirred for an additional 1h at –78 °C before being allowed to warm to room temperature overnight (ca. 12 h). The reaction mixture was then concentrated under reduced pressure and the residue was by purified column chromatography (SiO_2_, 10% EtOAc-hexanes) to provide **6** as a colorless oil (49 mg, 42%). ^1^H NMR (CDC1_3_, 400 MHz) δ: 5.34 (m, 2H), 3.66 (s, 3H), 2.30 (t, *J* = 7.5 Hz, 2H), 2.18 (m, 2H), 2.01 (m, 4H), 1.93 (t, *J* = 2.6 Hz, 1H), 1.61 (m, 4H), 1.51 (m, 4H), 1.26 (m, 18H). LRMS-ESI *m/z* calculated for C_23_H_41_O_2_^+^ [M+H^+^] calculated: 349.31, found 349.31

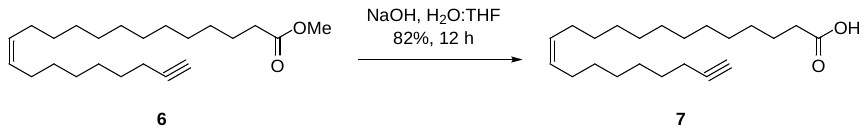

**(*Z*)-Docos-13-en-21-ynoic Acid (7)**: Methyl ester **6** (140 mg, 0.46 mmol, 1.0 equiv) was dissolved in THF (1.0 mL), and NaOH (91 mg, 2.28 mmol, 5.0 equiv) in H_2_O (1.0 mL) was added. The reaction mixture was stirred at room temperature for 8 h and then transferred to a separatory funnel containing EtOAc (25 mL) and aqueous 0.1 M HCl (25 mL). The product was extracted with EtOAc (3 x 25 mL), and the combined organic layers were dried over anhydrous Na_2_SO_4_ and concentrated under reduced pressure. The residue was purified by column chromatography (SiO_2_, 30% EtOAc-hexanes) providing **7** as a colorless oil (110 mg, 82%). ^1^H NMR (CDC1_3_, 400 MHz) δ: 5.34 (m, 2H), 2.30 (t, *J* = 7.5 Hz, 2H), 2.18 (m, 2H), 2.01 (m, 4H), 1.93 (t, *J* = 2.6 Hz, 1H), 1.61 (m, 4H), 1.51 (m, 4H), 1.26 (m, 18H). LRMS-ESI *m/z* calculated for C_22_H_39_O_2_^+^ [M+H^+^] calculated: 335.30, found 335.30.

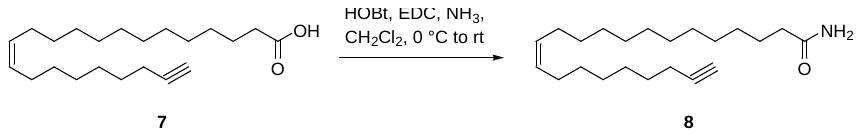

**(*Z*)-Docos-13-en-21-ynamide (8)**: Carboxylic acid **7** (42 mg, 0.12 mmol, 1 equiv) was dissolved in CH_2_Cl_2_ (2.5 mL) and the solution was cooled to 0 °C. HOBt (48.6 mg, 0.36 mmol, 3 equiv) was added followed by EDC (69 mg, 0.36 mmol, 3 equiv) and the reaction mixture was stirred for 30 minutes at 0 °C and 30 minutes at room temperature. The reaction mixture was cooled to 0 °C and 0.5 M NH_3_^.^in THF (1.2 mL, 0.60 mmol, 5 equiv) was added and the mixture slowly warmed to room temperature and stirred for an additional 30 minutes. The reaction mixture was diluted with CH_2_Cl_2_, washed with water, dried over Na_2_SO_4_, filtered and concentrated. Column chromatography (SiO_2_, 70% EtOAc-hexanes) yielded **8** as a white solid (33.6 mg, 81%).^1^H NMR (CDC1_3_, 400 MHz) δ: 5.49-5.25 (m, 4H), 2.27-2.14 (m, 4H), 2.01 (m, 4H), 1.93 (t, *J* = 2.6 Hz, 1H), 1.61 (m, 5H), 1.51 (m, 2H), 1.26 (m, 19H). LRMS-ESI *m/z* calculated for C_22_H_40_NO^+^ [M+H^+^] calculated: 333.30, found 333.30.

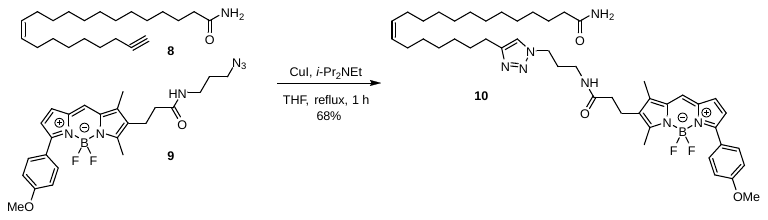

**^BDP^Eru** **(10)**: A solution of amide **8** (4.6 mg, 0.014 mmol, 1.3 equiv) in THF (0.5 mL) was treated with BDP-azide **9** (5 mg, 0.01 mmol, 1 equiv) followed by CuI (0.95 mg, 0.005 mmol, 0.5 equiv) and *i*-Pr_2_NEt (7 uL, 0.0035 mmol, 3.5 equiv) and warmed at reflux for 1 h. The reaction mixture was concentrated, and the residue was purified by preparative thin layer chromatography (10% MeOH-CH_2_Cl_2_) to yield **10** as a pink solid (5.6 mg, 68%). ^1^H NMR (CDC1_3_, 400 MHz) δ: 7.87 (d, *J* = 8.7 Hz, 2H), 7.06 (s, 1H), 6.96 (d, *J* = 8.7 Hz, 2H), 6.94 (d, *J* = 3.9 Hz, 1H), 6.01 (t, *J* = 5.7 Hz, 1H), 5.54-5.29 (m, 4H), 4.20 (t, *J* = 6.3 Hz, 2H), 3.85 (s, 3H), 3.17 (q, *J* = 5.9 Hz, 2H), 2.77 (t, *J* = 7.5 Hz, 2H), 2.64 (t, *J* = 7.5 Hz, 2H), 2.53 (s, 3H), 2.30 (t, *J* = 7.1 Hz, 2H), 2.22 (s, 3H), 2.20 (t, *J* = 7.5 Hz, 2H), 2.00 (s, 7H), 1.62 (m, 27H).

**Eru.da**

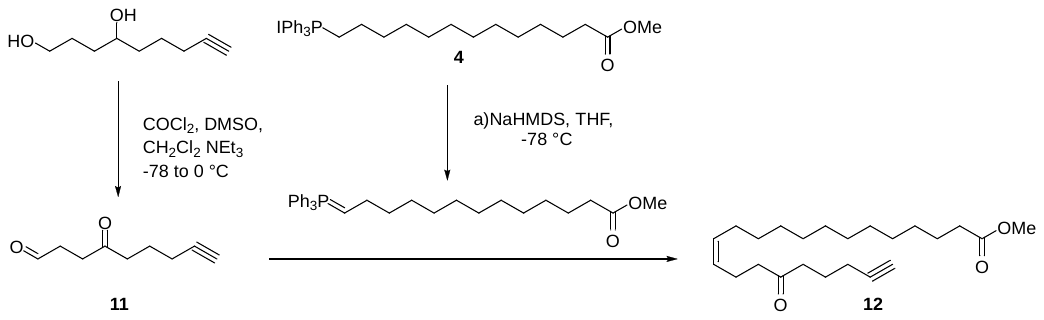

**Methyl (*Z*)-17-Oxodocos-13-en-21-ynoate (12)**: A stirred solution of phosphonium salt **4** (208 mg, 0.34 mmol, 1.0 equiv) in anhydrous THF (15 mL) at –78 °C under N_2_ was treated with NaHMDS (0.47 mL, 0.47 mmol, 1.0 M in THF, 1.0 equiv) dropwise over 15 min. The resulting orange solution was stirred at –78 °C for an additional 1 h. In a separate round-bottomed flask, aldehyde **11**^S1^ (86 mg, 0.56 mmol, 1.2 equiv) was dissolved in anhydrous THF (15 mL) and cooled to –78 °C under N_2_. The ylide solution was then slowly added to the aldehyde solution over 45 min and stirred for an additional 1 h at –78 °C, before the reaction mixture was allowed to warm to room temperature overnight (ca. 12 h). The reaction was then concentrated under reduced pressure and the residue was purified by column chromatography (SiO_2_, 10% EtOAc-hexanes) to provide **12** as a colorless oil (51 mg, 42%). ^1^H NMR (CDCl_3_, 500 MHz) δ: 5.46-5.26 (m, 2H), 3.69 (s, 3H), 2.58 (t, *J* = 7.2 Hz, 2H), 2.48 (m, 2H), 2.32 (m, 4H), 2.25 (dt, *J* = 7.0, 2.5 Hz, 2H), 2.05 (q, *J* = 6.8 Hz, 2H), 1.95 (t, *J* = 2.6 Hz, 1H), 1.83 (p, *J* = 7.2 Hz, 2H), 1.64 (m, 2H), 1.28 (m, 16H). LRMS-ESI *m/z* calculated for C_23_H_39_O_3_^+^ [M+H^+^] calculated: 363.28, found 362.27.

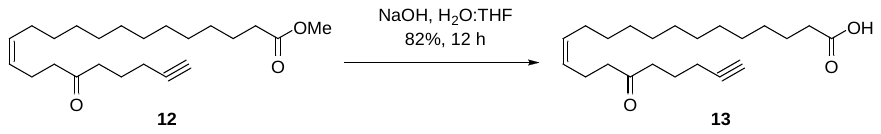

**(*Z*)-17-Oxodocos-13-en-21-ynoic Acid (13)**: Methyl ester **12** (166 mg, 0.46 mmol, 1.0 equiv) was dissolved in THF (1.0 mL), and NaOH (91 mg, 2.28 mmol, 5.0 equiv) in H_2_O (1.0 mL) was added. The reaction mixture was stirred at room temperature for 8 h and then transferred to a separatory funnel containing EtOAc (25 mL) and aqueous 0.1 M HCl (25 mL). The product was extracted with EtOAc (3 x 25 mL), and the combined organic layers were dried over anhydrous Na_2_SO_4_ and concentrated under reduced pressure. The residue was purified by column chromatography (SiO_2_, 30% EtOAc-hexanes) to provide **13** as a colorless oil (131 mg, 82%). ^1^H NMR (CDCl_3_, 500 MHz) δ: 5.46-5.26 (m, 2H), 2.55 (t, *J* = 7.2 Hz, 2H), 2.46 (t, *J* = 7.4 Hz, 2H), 2.32 (m, 4H), 2.25 (dt, *J* = 7.0, 2.5 Hz, 2H), 2.05 (q, *J* = 6.6 Hz, 2H), 1.95 (t, *J* = 2.5 Hz, 1H), 1.79 (p, *J* = 7.2 Hz, 2H), 1.64 (m, 2H), 1.28 (m, 16H). LRMS-ESI *m/z* calculated for C_22_H_37_O_3_^+^ [M+H^+^] calculated: 349.27, found 349.28.

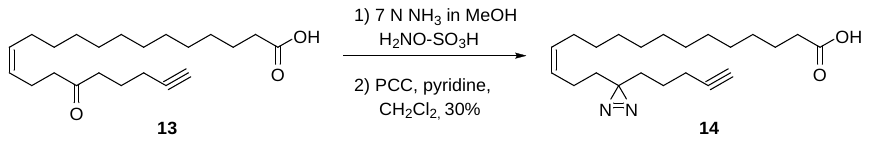

**(*Z*)-16-(3-(Pent-4-yn-1-yl)-3*H*-diazirin-3-yl)hexadec-13-enoic Acid** **(14)**: Ketone **13** (88 mg, 0.25 mmol, 1.0 equiv) was dissolved in a solution of NH_3_ in MeOH (2.0 mL, 7.0 N in MeOH) at 0 °C in a round-bottom flask under N_2_. After stirring for 3 h at 0 °C, a solution of hydroxylamine-*O*-sulfonic acid (32.9 mg, 0.29 mmol, 1.15 equiv) in anhydrous MeOH (2.0 mL) was added dropwise over 10 min. The reaction mixture was allowed to slowly warm to room temperature while stirring overnight (ca. 12 h). The reaction mixture was then concentrated under a stream of N_2_ and the residue was redissolved in Et_2_O (10 mL) and filtered through a pad of Celite using additional Et_2_O (5 mL) for washes. The combined filtrate was concentrated under reduced pressure providing the crude diaziridine intermediate which was used without further purification.

The crude diaziridine intermediate was dissolved in anhydrous CH_2_Cl_2_ (4.0 mL) and pyridine (0.4 mL) in a round-bottom flask charged with a stir bar and flushed with N_2_. Pyridinium chlorochromate (109 mg, 0.51 mmol, 2.0 equiv) was then added in small portions over 20 min while the reaction mixture was cooled at 0 °C. The reaction mixture was then allowed to warm to room temperature and stirred for an additional 1 h before diluting with 50% EtOAc/hexanes (25 mL). The resulting solution was passed through a silica gel plug and the filtrate was concentrated and purified further by flash chromatography (SiO_2_, 20% EtOAc-hexanes,) to provide **14** as a colorless oil (27.6 mg, 30%, 2 steps). ^1^H NMR (CDCl_3_, 500 MHz) δ: 5.37 (m, 1H), 5.23 (m, 1H), 2.34 (t, *J* = 7.3 Hz, 2H), 2.16 (dt, *J* = 6.6, 2.6 Hz, 2H), 1.98 (q, *J* = 7.1 Hz, 2H), 1.94 (t, *J* = 2.7 Hz, 1H), 1.83 (q, *J* = 7.2 Hz, 2H), 1.63 (p, *J* = 7.5 Hz, 2H), 1.51 (m, 2H), 1.43 (m, 2H), 1.37-1.20 (m, 18H). LRMS-ESI *m/z* calculated for C_22_H_37_N_2_O_2_^+^ [M+H^+^] calculated: 361.28, found 361.28.

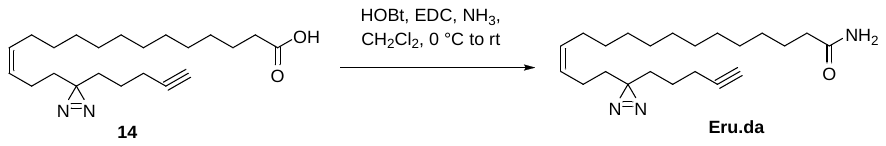

**(*Z*)-16-(3-(Pent-4-yn-1-yl)-3*H*-diazirin-3-yl)hexadec-13-enamide** **(Eru.da)**: Carboxylic acid **14** (27.8 mg, 0.08 mmol, 1 equiv) was dissolved in CH_2_Cl_2_ (2.5 mL) and cooled to 0 °C. HOBt (31.2 mg, 0.23 mmol, 3 equiv) was added followed by EDC (44.2 mg, 0.23 mmol, 3 equiv) and the mixture was stirred for 30 min at 0 °C and 30 minutes at room temperature. The reaction mixture was cooled to 0 °C and 0.5 M NH_3_ in THF (0.77 mL, 0.38 mmol, 5 equiv) was added. The mixture was slowly warmed to room temperature and stirred for an additional 30 minutes. The reaction mixture was diluted with CH_2_Cl_2_, washed with water, dried over Na_2_SO_4_, filtered and concentrated. Column chromatography (SiO_2_, 1% MeOH-EtOAc) yielded **Eru.da** as a white solid (19.5 mg, 71%).^1^H NMR (CDCl_3_, 500 MHz) δ: 5.47-5.19 (m, 4H), 2.22 (t, *J* = 7.5 Hz, 2H), 2.16 (td, *J* = 7.8, 2.5 Hz, 2H), 1.98 (q, *J* = 6.8 Hz, 2H), 1.95 (t, *J* = 2.6 Hz, 1H), 1.83 (q, *J* = 7.2 Hz, 2H), 1.64 (q, *J* = 7.1 Hz, 2H), 1.51 (m, 2H), 1.43 (m, 2H), 1.31 (m, 18H). ^13^C NMR (126 MHz, CDCl_3_) δ: ^13^C NMR (126 MHz, CDCl_3_) δ 175.8, 131.4, 127.8, 83.6, 69.0, 36.1, 33.2, 31.9, 29.70, 29.68, 29.64, 29.59, 29.46, 29.41, 29.36, 28.5, 27.3, 25.7, 22.9, 21.8, 18.1 (δ of one carbon absent). HRMS-ESI *m/z* calculated for C_22_H_38_N_3_O^+^ [M+H^+^] calculated: 360.3010, found 360.3009.

**Ole.da**

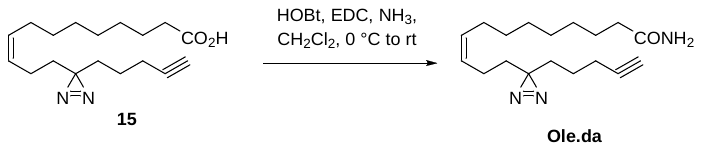

**(*Z*)-12-(3-(Pent-4-yn-1-yl)-3*H*-diazirin-3-yl)dodec-9-enamide** **(Ole.da)**: Carboxylic acid **15** (25 mg, 0.08 mmol, 1 equiv) was dissolved in CH_2_Cl_2_ (2.5 mL) and cooled to 0 °C. HOBt (33.7 mg, 0.25 mmol, 3 equiv) was added followed by EDC (47.9 mg, 0.25 mmol, 3 equiv). The mixture was stirred for 30 minutes at 0 °C and 30 min at room temperature. The reaction mixture was cooled to 0 °C and 0.5 M NH_3_ in THF (0.8 mL, 0.60 mmol, 5 equiv) was added and the mixture slowly warmed to room temperature and stirred for an additional 30 min. The reaction mixture was diluted with CH_2_Cl_2_, washed with water, dried over Na_2_SO_4_, filtered and concentrated. Column chromatography (SiO_2_, 1% MeOH-EtOAc) yielded **Ole.da** as a white solid (10 mg, 42%).^1^ ^1^H NMR (CDCl_3_, 500 MHz) δ: 5.44 (m, 2H), 5.36 (m, 1H), 5.24 (m, 1H), 2.22 (t, *J* = 7.5 Hz, 2H), 2.16 (td, *J* = 7.0, 2.5 Hz, 2H), 1.98 (q, *J* = 6.2 Hz, 2H), 1.95 (t, *J* = 2.8 Hz, 1H), 1.82 (q, *J* = 7.6 Hz, 2H), 1.64 (q, *J* = 7.1 Hz, 2H), 1.51 (m, 2H), 1.43 (m, 2H), 1.31 (m, 10H). ^13^C NMR (126 MHz, CDCl_3_) δ 131.3, 127.9, 90.1, 83.6, 69.1, 36.1, 33.2, 31.0, 29.6, 29.33, 29.31, 29.2, 28.5, 25.7, 22.9, 21.8, 18.1 (δ of one carbon absent). HRMS-ESI *m/z* calculated for C_18_H_30_N_3_O^+^ [M+H^+^] calculated: 304.2304, found 304.2302.

**Pal.da**

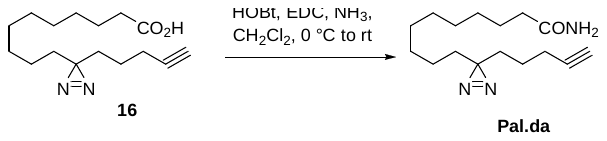

**10-(3-(Pent-4-yn-1-yl)-3*H*-diazirin-3-yl)decanamide (Pal.da)**: Carboxylic acid **16^1^** (23 mg, 0.08 mmol, 1 equiv) was dissolved in CH_2_Cl_2_ (2.5 mL) and cooled to 0 °C. HOBt (33.7 mg, 0.25 mmol, 3 equiv) was added followed by EDC (47.9 mg, 0.25 mmol, 3 equiv) and the mixture was stirred for 30 minutes at 0 °C and 30 minutes at room temperature. The reaction mixture was cooled to 0 °C and 0.5 M NH_3_ in THF (0.8 mL, 0.41 mmol, 5 equiv) was added and the mixture was slowly warmed to room temperature and stirred for an additional 30 minutes. The reaction mixture was diluted with CH_2_Cl_2_, washed with water, dried over Na_2_SO_4_, filtered and concentrated. Column chromatography (SiO_2_, 80% EtOAc-hexanes) yielded **Pal.da** as a white solid (12.5 mg, 55%). ^1^H NMR (CDCl_3_, 500 MHz) δ: 5.94 (m, 2H), 2.21 (t, *J* = 7.6 Hz, 2H), 2.16 (dt, *J* = 7.0, 2.5 Hz, 2H), 1.94 (t, *J* = 2.6 Hz, 1H), 1.62 (q, *J* = 7.1 Hz, 2H), 1.48 (m, 2H), 1.39-1.14 (m, 16H). HRMS-ESI *m/z* calculated for C_16_H_28_N_3_O^+^ [M+H^+^] calculated: 278.2227, found 278.2227.

**The gating strategy for FAC sorted CD11b+ cells in Fig. 2e**

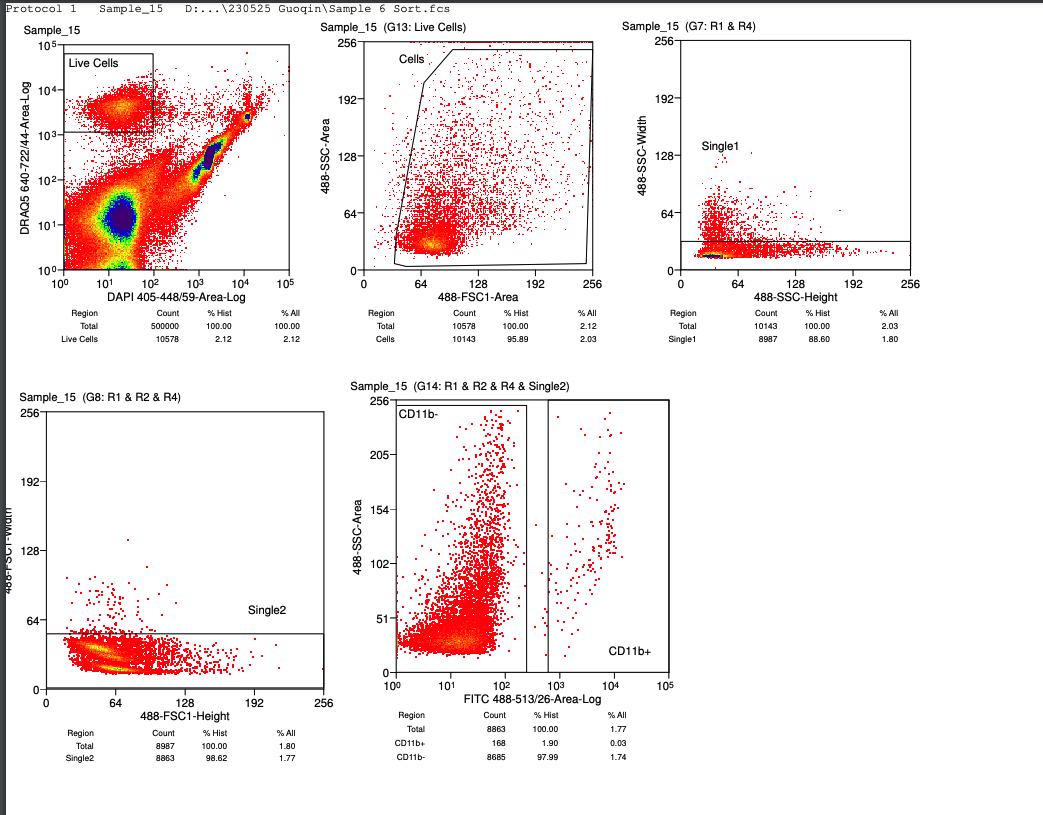

Step 1: Sort Plots as indicated in the first box of “Live cells” DAPI vs DRAQ5. DRAQ5 separates cells from debris.  DAPI is used to distinguish live cells from dead.

Step 2: Scatter Plots as indicated in the box of “Cells” FSC vs SSC 488nm forward scatter (FSC) information on cell size.  488nm side scatter (SSC) information on cell shape or complexity. Used to distinguish cell types and separates cells from debris.

Step 3: Doublet Discrimination Plots as indicated in the box of “Single1” and “Single2” SSC Pulse Height vs SSC Pulse Width & FSC Pulse Height vs FSC Pulse Width Used to exclude doublets from the sorted population.

Step 4: Fluorescence Plot as indicated in the box of “CD11b+” FITC CD11b vs SSA Used to distinguish CD11b positive and negative cells. CD11b+ cells were used for RNA extraction and qPCR in Fig. 2e.

**References**

1.	Niphakis, M.J.*, et al.* A Global Map of Lipid-Binding Proteins and Their Ligandability in Cells. *Cell* **161**, 1668-1680 (2015).
